# Prefrontal deep projection neurons enable cognitive flexibility via persistent feedback monitoring

**DOI:** 10.1101/828590

**Authors:** Spellman Timothy, Svei Malka, Kaminsky Jesse, Manzano-Nieves Gabriela, Liston Conor

## Abstract

Cognitive flexibility, the ability to alter one’s strategy according to changing stimulus-response-reward relationships, is critical for acquiring and updating learned behavior. Attentional set-shifting, a test of cognitive flexibility, depends on the activity of prefrontal cortex (PFC). It remains unclear, however, what specific role PFC neurons play and how they interact to support set-shifting. One widely held view is that prefrontal activity biases sensorimotor responses by mediating attention. Using optogenetics and 2-photon calcium imaging, we demonstrate that, while PFC activity does encode attentional sets, this activity does not bias sensorimotor responses. Rather, PFC activity enables set-shifting by encoding trial feedback information, a role it has been known to play in other contexts. We identify a circuit-level mechanism that supports feedback monitoring through persistent, recurring activity bridging multiple trials. Unexpectedly, the functional properties of PFC cells did not vary with their efferent projection targets in this context. Instead, representations of trial feedback formed a topological gradient, with cells more strongly selective for feedback information located further from the pial surface and receiving denser afferent inputs from the anterior cingulate cortex. Together, these findings identify a critical role for deep PFC projection neurons in enabling set-shifting through behavioral feedback monitoring.

## Introduction

*Cognitive flexibility,* the ability to respond to changing environmental contingencies, is critical for any organism that must navigate through a dynamic environment. A form of cognitive flexibility frequently employed in clinical assessments is *attentional set-shifting*, a kind of task-switching behavior that requires a subject to ignore a previously relevant stimulus feature and instead attend to a previously irrelevant feature (Heisler et al., 2015; Tait et al., 2014).

Set-shifting impairments are common in a range of psychiatric disorders (Ceaser et al., 2008; Disner et al., 2011; Halleland et al., 2012; Jazbec et al., 2007; Murphy et al., 2012) and often persist after treatment in both depression and schizophrenia, despite remission of other symptoms (Bortolato et al., 2016; Gonda et al., 2015; Harvey et al., 2004). A long-standing body of evidence indicates that the prefrontal cortex (PFC) plays a critical role in supporting set-shifting behavior in human (Kim et al., 2012; Milner, 1963) and rodent models (Birrell and Brown, 2000; Bissonette et al., 2013; Brigman et al., 2005), but the circuit elements and physiological mechanisms that enable set-shifting remain poorly resolved. A functional dissection of the prefrontal circuitry underlying set-shifting is of particular translational interest because pharmacotherapeutic interventions in this circuitry can have complex or paradoxical effects by acting on competing circuit elements (Miller, 2016), which may account for the deleterious cognitive side effects that can accompany antipsychotics and SSRIs (Hill et al., 2010; Sayyah et al., 2016).

The dominant model for the role of PFC in set-shifting, which has drawn on support from numerous high-impact publications over the past three decades (Miller and Cohen, 2001; MacDonald et al., 2000; Corbetta and Shulman, 2002; Desimone and Duncan, 1995; Birrell and Brown, 2000; Schmitt et al., 2017), holds that the PFC supports set-shifting by encoding abstract task rules and attentional sets that mediate top-down control of sensorimotor processing and decision making. There is physiological support for this model from animal studies. Neural activity in primate prefrontal cortex encodes abstract contextual or rule-related information (Hymana et al., 2012; Mante et al., 2013; Rigotti et al., 2013), and in rodents performing set-shifting tasks, such rule-related representations can shift flexibly with repeatedly changing stimulus-reward contingencies (Durstewitz et al., 2010; Mante et al., 2013; Rich and Shapiro, 2009; Rodgers and DeWeese, 2014; Siniscalchi et al., 2016). Together, these studies suggest that PFC activity might provide an attentional filter that biases sensorimotor responses during set-shifting (Wimmer et al., 2015)—a well-predicated but as yet unproven hypothesis.

Importantly, set-shifting tasks are typically uncued, mirroring the need for uncued adaptations to changing environmental contingencies in the real world, so set-shifting performance requires continuous trial-and-error learning. Thus, an alternative and not mutually exclusive hypothesis is that PFC supports set-shifting by monitoring feedback in response to recent decisions. In addition to encoding context, prefrontal activity has been shown to represent feedback signals associated with trial outcomes (Bissonette and Roesch, 2015; Starkweather et al., 2018), and recent evidence suggests that such feedback-related activity is important for task-switching behavior (Biró et al., 2019; Ellwood et al., 2017). Whether PFC activity supports set-shifting through feedback monitoring or through attentional modulation of sensorimotor responses is unknown.

It is also unclear how distinct circuit elements interact within PFC to support set-shifting. A major set of questions in behavioral neuroscience in recent years has centered on how best to define anatomical markers of cell types within cortex. Are the functional/coding properties of a given pyramidal neuron determined by its *efferent connectivity* or by its laminar location, a correlate of its *afferent connectivity* (Adesnik and Naka, 2018; Harris and Shepherd, 2015). These two possibilities are not mutually exclusive, and the fact that neuronal subpopulations with distinct efferent connectivity profiles are often found in distinct cortical layers makes it hard to disambiguate the relative contributions of these two factors. While findings from numerous, recent studies implicate both laminar location and efferent connectivity in determining the functional properties of pyramidal neurons (Lui et al., 2020; Marshel et al., 2019; Senzai et al., 2019; Sharif et al., 2020), to disambiguate these two factors requires an experimental preparation that controls for both. We therefore employed a virally-mediated expression approach that allowed for targeted labeling of PFC cell types that differed according to their projection targets or laminar location.

The critical contribution of PFC activity to set-shifting may be mediated by a number of output pathways, but two PFC projection targets of particular interest are the projection to ventromedial striatum (PFC-VMS) and to the medio-dorsal thalamus (PFC-MDT). In rodents, both target structures have established roles in set-shifting behavior (Aoki et al., 2015; Block et al., 2007; Floresco et al., 2006; Kato et al., 2018), and PFC projections to both structures underlie behavior that relies on cognitive flexibility (Marton et al., 2018; Nakayama et al., 2018). However, it is unknown whether PFC projections to these downstream targets convey similar or distinct task-critical information during set-shifting.

To address these questions, we developed and validated a novel set-shifting task for head-fixed mice to enable 2-photon imaging during serial attentional set shifts spanning hundreds of trials. While PFC neural activity encoded all essential task features, neural signals encoding the animal’s response were detected only *after* trial completion, supporting a role for PFC cells in feedback monitoring rather than attentional biasing of sensorimotor responses. This feedback signal was persistent and detectable through four subsequent trials, spanning up to 55 seconds. Separate analysis of PFC-VMS and PFC-MDT neurons revealed strikingly similar representations of all task-related features in both cell types. Unexpectedly, whereas optogenetic inhibition of either cell type had no effect on performance when delivered *during trials*, inhibition during the *post-trial epoch* did impair performance—but only following rule-dependent trials, confirming that the role of these neurons in set-shifting was feedback monitoring and not attentional modulation of sensorimotor responses. In contrast with PFC, inhibition of posterior parietal cortex (PPC) *during trials* did impair performance, showing that PPC but not PFC is required for attentional modulation of sensorimotor responses. Furthermore, while the functional properties of PFC cells did not vary with their efferent projection targets in this context, we found that representations of trial feedback formed a topological gradient, with cells more strongly selective for feedback information located further from the pial surface and receiving denser afferent inputs from the caudo-ventral anterior cingulate cortex (ACC). Together, these findings reveal a critical role for deep PFC projection neurons in supporting set-shifting by relaying feedback information to downstream targets.

## Results

### A cross-modal set-shifting task for head-fixed mice

To image PFC activity during set-shifting behavior in mice, we developed a head-fixed paradigm leveraging the sensitivity of the olfactory and whisker somatosensory systems and permitting us to record from hundreds of cells during serial attentional set shifts spanning hundreds of trials. Water-restricted mice were presented on each trial with one of two possible whisker vibration stimuli (e.g. 35Hz vs 155Hz vibration presented bilaterally) and one of two possible odor stimuli (e.g. almond oil vs olive oil), in randomized combinations, cueing them to respond by licking a left or right lick ports to retrieve a water reward upon termination of the 2.5s compound stimulus (**Fig. 1A-B**). As in most set-shifting tasks (Bissonette et al., 2008; Tait et al., 2014), animals underwent a standardized series of task transitions in order to expose them to multiple exemplars from each stimulus modality and build an attentional set: simple discrimination (SD), in which animals were trained to discriminate between two stimuli within a single sensory modality; compound discrimination (CD), in which a distractor stimulus from the untrained sensory modality was added; intradimensional shift (IDS), in which the stimuli from the relevant sensory modality were replaced with a new pair of exemplars; reversal (Rev), in which the left/right mapping was switched; extradimensional shift (EDS), in which the rule changed for the first time so that the pair of stimuli from the previously *irrelevant* sensory modality became the *relevant* stimuli; a second intradimensional shift (IDS2), in which the pair of stimuli from the *newly relevant* sensory modality was replaced with a new pair of exemplars; and finally serial extradimensional shifts (SEDS), in which the rule switched automatically whenever the animal reached criterion performance, which was 80% correct within a 30-trial moving window, and >50% on both left and right trials (**Fig. 1A,** see Methods for details). To validate the set-shifting task, we confirmed that animals took longer to perform an EDS than an IDS, and there was no difference between modalities in trials to criterion (**Fig. 1C**). Infusion of muscimol within PFC (**Fig. 1D**) impaired set-shifting performance, increasing trials to criterion in both the initial EDS shift, as well as after completing 10 shifts of SEDS (**Fig. 1E**), showing that set-shifting in this paradigm is dependent on PFC activity, both initially and after repeated shifts.

**Figure 1:**
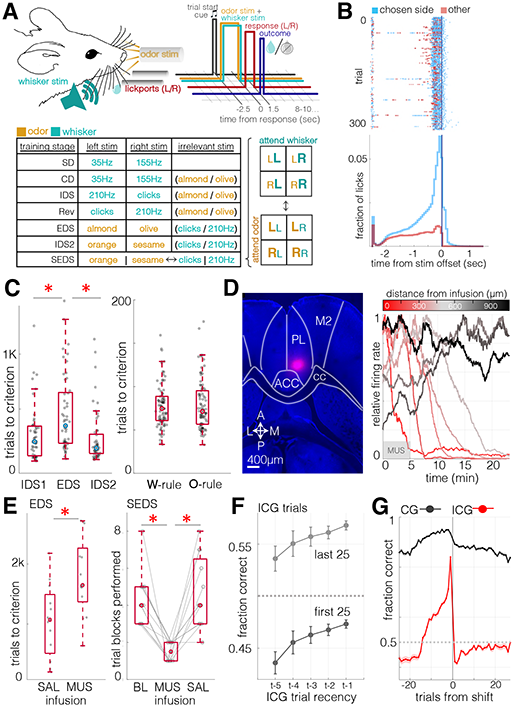
A prefrontal-dependent serial set-shifting task for head-fixed mice. A. Task schematic. Top left: diagram of stimulus delivery and lick response configuration. Top right: trial sequence. A trial begins with a 500ms white-noise trial-start cue, followed by a 2.5s presentation of whisker and/or odor stimulus. Following stimulus offset, there is a response window (≤1.5s) during which a lick to either lick port terminates the trial, and a correct response is rewarded with a 3μL water bolus. Bottom: a table of the discrete training stages culminating in the serial extra-dimensional set-shifting task. <u>SD</u> = Simple Discrimination (single stimulus modality), <u>CD</u> = Compound Discrimination (a second, irrelevant stimulus modality is added), <u>IDS1</u> = Intradimensional Shift (stimuli within the initially relevant modality are replaced with a new set), Rev = Reversal (left/right stimulus/reponse mapping is reversed within initial modality rule), <u>EDS</u> = Extradimensional Shift (same stimuli, new modality rule), <u>IDS2</u> = Intradimensional Shift (stimuli within the newly relevant modality are replaced with a new set), <u>SEDS</u> = Serial Extradimensional Shifting (the modality rule is repeatedly switched). B. Summary of licking behavior. Top: rasterized and trial-aligned lick times for an example session. Licks toward the ultimately chosen lick port in blue, licks toward the other lick port in red. On many trials, both lick ports are explored prior to the response window. Bottom: Summary lick time histogram (655,236 licks, 150 sessions, 32 animals). On 95% of trials, the choice lick (first lick made after stimulus termination) came in the first 0.346ms of the response window, a latency equal to a single frame of imaging. C. Left: Number of trials to criterion, Intradimensional vs Extradimensional Shift, n=53 animals. Signed rank z = −2.9, p = 0.0034, for IDS1/EDS; z = −4.0, p = 6× 10^−5^ for IDS2/EDS, z = 1.4, p = 0.15 for IDS1/IDS2. Right: Mean trials to criterion during SEDS sessions, whisker rule vs odor rule. N=115 animals, signed rank z = 0.14, p = 0.9. D. Muscimol infusion in PFC. Left: An infusion of fluorescent muscimol made using same concentration, volume, and infusion rate as in the muscimol behavioral experiment. Right: Relative multiunit firing rate (proportion of channel maximum, 0.5Hz bins, 100s moving average). Each trace is the mean activity of a pair of electrodes at a given A/P distance from the infusion site. E. Left: trials to criterion in EDS sessions during transcranial infusion (SAL = saline; MUS = muscimol). N= 12, 13 mice (SAL, MUS). Rank sum z = 2.4, p = 0.02. Right: Number of trial blocks reaching criterion performance in SEDS sessions following 10 rule shifts. N=12; median blocks (BL/MUS/SAL): 4, 1.5, 4. Signed rank p = 0.0005 (BL/MUS), 0.001 (SAL/MUS), 0.68 (BL/SAL). Median total trials completed: SAL = 644; MUS = 651; Signed rank p = 0.52. F. Incongruent trial performance by recency of previous incongruent trial during SEDS sessions (mean ± SEM). Top: performance during the last 25 trials of trial blocks, prior to reaching criterion; bottom: performance during the first 25 trials of trial blocks, after undergoing rule shifts. N = 693 sessions in 131 animals. ANOVA for ICG trial recency vs ICG trial performance: t = −4.68, p = 3× 10^−6^. G. SEDS trial performance relative to rule shift, mean ± SEM. Same sessions as in **F**.

Trials belonged to two classes: those in which the whisker and odor rules cued the same response direction (*congruent* trials: CG), and those in which the whisker and odor rules cued opposing response directions (*incongruent* trials: ICG). Therefore, ICG trials, but not CG trials, required application of the modality rule, and feedback from ICG trials alone carried information about modality rule. Animals used prior ICG trial information to guide behavior, so that ICG trial performance was enhanced following feedback from a recent ICG trial, and this held true in trials occurring both early and late within trial blocks (**Fig. 1F**). Performance on ICG trials, but not CG trials, dropped below chance immediately following rule switches, revealing a perseverance of the previous rule; sub-chance performance persisted over multiple ICG trials, gradually reaching chance and then above-chance performance as new the new rule was acquired (**Fig. 1G**).

### Set-shifting task variables represented in prefrontal population activity

To measure task-related activity in PFC neurons, GCaMP6f-mediated 2-photon calcium imaging was performed through a coronally-implanted microprism (Andermann et al., 2013; Low et al., 2014), producing a field of view that preserved cortical laminar structure and was restricted to prelimbic and infralimbic areas (**Fig. 2A-B**). This field of view allowed for monitoring of a 1500μm × 760μm area, with a median of 450 active neurons per animal in a given session (registering cells across consecutive sessions yielded a median of 179 per animal – see Methods and **Fig. S3–S10**). GCaMP was expressed pan-neuronally (hSyn-GCaMP6f) or in specific projection neuron subtypes (hSyn-DIO-GCaMP6f and rAAV2-Cre in PFC-VMS and PFC-MDT). Although these cell types were subsequently analyzed separately, we began by characterizing the task-responsiveness of all labeled neurons. Examination of activity traces averaged over feature-matched trials revealed neurons whose response profiles displayed compound selectivity for multiple task features, such as whisker stimulus and odor stimulus (**Fig. 2C**, left).

**Figure 2.**
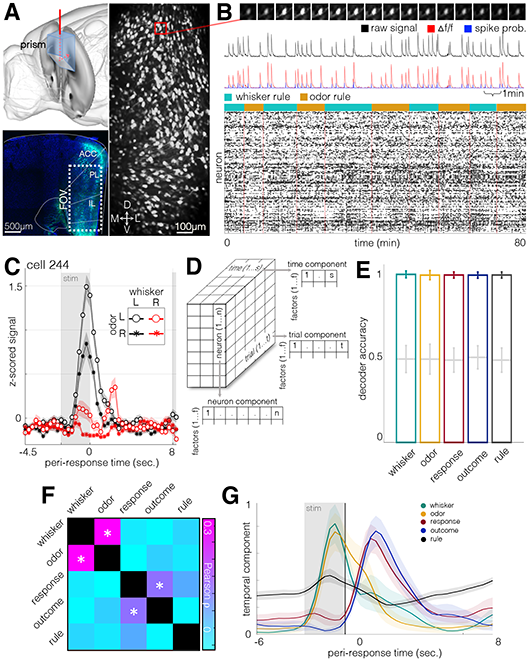
Prefrontal representation of task variables in population activity. A. Top left: schematic of coronal prism implantation field. FOV bregma coordinates: AP +1.85mm, ML 0-0.75mm left, DV 1.0mm-2.5mm from brain surface. Bottom left: hSyn-GCaMP6f fluorescence in fixed tissue (DAPI in blue). Right: standard deviation image from 115-minute hSyn-GCaMP6f recording (2.89Hz, N = 953 units). Note: different animal from bottom left. B. Example frames, traces, and putative spike times from calcium imaging sessions. Top: 16 frames of a calcium transient from an example neuron in (A). Frames are downsampled at a ratio of 1:2, so that the sequence shown is at 1.45Hz. Black trace: raw image fluorescence. Red trace: denoised △f/f. Blue: putative spike times from deconvolved △f/f signal. Bottom: rasterized putative spike times from SEDS session. N=100 units recorded over the course of 10 trial blocks. C. Mean ± SEM of z-scored activity for all trials from an example neuron during trials with four stimulus combinations (*whisker stimulus* right/left, *odor stimulus* right/left). Stimulus is presented from −2.5s to 0s. Earliest subsequent trial onset at 8 sec. D. Schematic of TCA rank decomposition. A three-dimensional data set (time of length s, neurons of length n, and trials of length t) is decomposed into three lower-rank components of f factors (s*f, n*f, and t*f). E. SVM decoder accuracy for five task-related features. Trial factors obtained from TCA decomposition (35 factors) were used as inputs to a support vector machine-based decoder, which was trained to classify trials according to five features: whisker stimulus, odor stimulus, response direction (left / right), trial outcome (correct / incorrect), and task rule (whisker rule / odor rule). Cross-validation was performed on trials (50%) held out from training. N=4740 neurons from 21 animals. Error bars are standard deviations. Gray lines and error bars are performance on shuffled labels. F. Cross-correlation of task feature representation in TCA trial space. GLM regression was performed on the five task variables, using TCA factors as predictors. Pearson correlation was then performed on the resulting coefficients (same data as in E). Asterisks indicate p<0.01. G. Time components from significantly modulated factors. For each of the five task variables, factors most strongly modulated by each task variable are shown in the temporal domain (mean +/− SEM for each variable, N=16 factors for whisker stimulus, N=12 factors for odor stimulus, N=26 factors for response direction, N=21 factors for trial outcome, N=59 factors for rule).

To ascertain which task-related features were encoded at the population level, we began by using tensor component analysis (TCA, **Fig. 2D**), to decompose the neural data into low-rank factors defined as related sets of weights in the neuron, trial, and timepoint (within-trial) dimensions (Williams et al., 2018). These weights reflect the magnitude of the underlying activity pattern at each neuron, trial, and timepoint. Because this dimensionality reduction technique separates trial and timepoint components, it allowed for variables to be directly compared with each other, regardless of when their peak representation occurred within the trial. Using the trial components as inputs to a support vector machine-based (SVM) maximum-margin linear decoder (Christianini and Shawe-Taylor, 2000; Meyers et al., 2008), trials were classified according to *whisker stimulus* (e.g. clicks vs. 210Hz), *odor stimulus* (e.g. almond vs. olive), *response* (left / right), *outcome* (correct / incorrect), and *rule* (whisker rule / odor rule) with near perfect accuracy in held-out test data (**Fig. 2E**).

We next sought to determine the degree to which representations of these task variables correlated with one another within the population activity space. To determine whether each task variable was encoded by a distinct pattern of activity from across the neuron population, or alternatively, whether multiple task variables were encoded by similar activity patterns, we performed general linear model (GLM) regression on the trial components, using the five variables as predictors, and then tested for correlations between the resulting coefficients across factors (**Fig. 2F**). Coefficients for *whisker* and *odor* stimuli were significantly correlated with each other across factors, as were *response* and *outcome*, indicating these pairs of variables were encoded by similar activity patterns. One possible way this could manifest is if activity patterns encoding whisker and odor follow overlapping temporal trajectories within the trial, and likewise for response and outcome. To determine the trial-aligned time-courses of the factors associated with the task variables, we plotted the TCA-derived temporal components of the factors most strongly associated with each variable (**Fig. 2G**). Temporal components associated with *whisker* and *odor* stimulus peaked during stimulus presentation, while components associated with *outcome* peaked during the subsequent inter-trial interval (ITI). Notably, components associated with *response* followed a temporal trajectory that was more similar to *outcome* than *whisker* or *odor* stimuli, peaking after the end of the trial (**Fig. 2G**). These results demonstrate that all task elements necessary for successfully executing the task are represented in PFC neuronal population activity, that activity patterns associated with *response* are more correlated with those associated with *outcome* than with those associated with the decision-related stimuli, and that the *response*-associated patterns lag, rather than lead, the animal’s behavioral choice.

### Response and outcome representations in post-trial activity

Next, we questioned how the activity of individual neurons contributed to the representation of task variables seen in the population-level analysis. Because the TCA-based population analysis revealed signals encoding *response* and *outcome* in the post-trial ITI, we wondered whether these variables might continue to be represented in the neural signal beyond the start of the subsequent trial. To answer this question, we used GLM regression to model activity rates in individual neurons as a function of *whisker stimulus, odor stimulus, response direction trial outcome, and rule*, with the addition of the variables *previous trial response* and *previous trial outcome*. **Fig. 3A-B** shows example trial-aligned activity traces, along with GLM estimates for trial-by-trial activity, for two example neurons encoding response and outcome in post-trial activity (see **Fig. S12** for further examples). Analysis of the mean GLM coefficients at each timepoint (**Fig. 3C**) revealed that coefficient values for *response* and *outcome* peaked at a mean of 1.73s and 2.42s after stimulus offset, respectively, and remained elevated well above values from shuffled data through the following trial.

**Figure 3:**
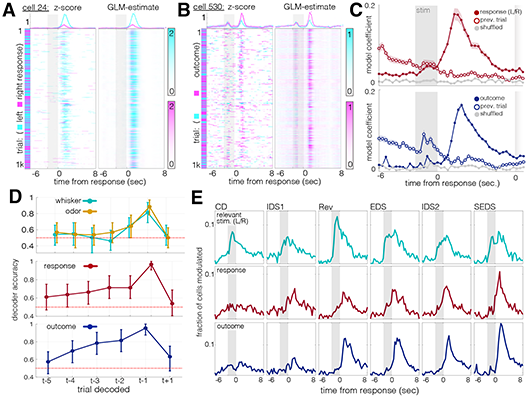
Task-related action and outcome representations in post-trial activity. A. Example neuron with selectivity for response direction in its post-trial activity. Left: z-scored activity. X-axis, time within trial (Gray box: stimulus presentation window, −2.5s – 0s). Y-axis: trial number (1071 trials total). Right: GLM estimate for time-aligned activity. Cyan: trials in which the animal chose left; magenta: trials in which the animal chose right. Top: mean +/− SEM traces from each trial condition. Top y-axis units are z-scored activity. B. Same format as (A), for an example neuron selective for trial outcome. Cyan: incorrect trials; magenta: correct trials. C. Trial-aligned GLM coefficients for neurons significantly modulated by response direction (top) and trial outcome (bottom). Modulated neurons include those with significant modulation at any timepoint (Bonferroni-corrected for the 43 trial timepoints). Closed circles: mean +/− SEM of coefficients for currently trial; open circles: mean +/− SEM of coefficients for previous trial. Gray traces: coefficients from GLMs on shuffled data. N=285 neurons for response direction, 756 for trial outcome, out of 4730 total neurons from 21 animals. D. Means ± 95% confidence intervals for SVM decoders tested on past (t−…) and future (t+…) trial features. Top: decoders for whisker and odor stimulus. Middle: decoders for response direction. Bottom: decoders for trial outcome. Red dashes: chance performance. E. Histograms: fraction of neurons significantly modulated (same GLM as A-C) by task features over trial-aligned timepoints and through learning stages. Top row: fraction of neurons modulated by the relevant stimulus. Middle row: response direction. Bottom row: trial outcome.

The observation that representations of *response* and *outcome* persisted through the subsequent trial raised the possibility that these signals might be detected beyond this time interval. Given that animals required 15-20 incongruent trials to abandon a rule after an uncued rule-change (**Fig. 1E**), we looked for evidence of outcome-related evidence accumulation that spanned multiple trials, as has been reported in reinforcement learning tasks (Bari et al., 2019; Bernacchia et al., 2011; Siniscalchi et al., 2019). We ran linear decoders on *whisker* and *odor stimuli*, *response*, and *outcome* for past trials and found that, while whisker and odor representations were no longer detectable after one trial, response and outcome representations persisted for up to four trials, or up to 55 seconds (**Fig. 3D**). Importantly, the inability to decode these features at above-chance accuracies from future trials (timepoint t+1 in **Fig 3D**) served as a negative control against potential autocorrelation effects in behavior and/or neural activity.

Do these *response* and *outcome* signals reflect the demands of the uncued set-shifting task, or are they natively expressed during decision-making more generally? To answer this question, we tracked response and outcome signals across learning stages. Signals corresponding with the relevant *stimulus* was present from the earliest recorded session (CD) and remained strong throughout all subsequent task stages, (**Fig. 3E**, top). Conversely, very few neurons were modulated by *response* or *outcome* in early (e.g. CD) sessions. Instead, *response-* and *outcome*-related activity emerged only over the course of multiple task transitions (**Fig. 3E,** rows 2 and 3). While this analysis does not precisely identify the minimum conditions necessary to evoke the *response* and *outcome* signals, the absence of these signals in early sessions means that these signals do not innately result from discrimination behavior, but require extensive training and/or task complexity.

### A circuit-level mechanism representing outcomes across multiple trials in a stable, colinear activity space

That trial feedback information could be decoded for up to four trials in the past led us to question how these past trial outcomes were encoded: were these representations stable, maintained by consistent groups of neurons, or by groups whose membership shifted over time? We first examined the stability of *outcome* selectivity across sessions. *Outcome* selectivity, defined hereafter as the difference between mean z-scored activity in correct and incorrect trials for each neuron, exhibited low correlation between pairs of early sessions, when the representations themselves were weak (**Fig. 4A**). Between later session pairs, however, selectivity became highly correlated, especially among SEDS sessions (**Fig. 4B-C**), indicating that outcome is represented by relatively stable groups of neurons over days.

**Figure 4:**
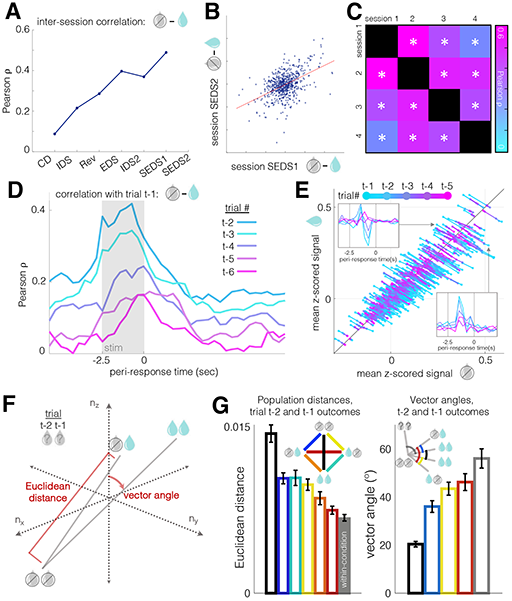
Outcomes are represented across multiple trials in a stable, colinear activity space. A. Inter-session correlation of trial outcome representation. X-axis: imaging sessions in which each significantly modulated neuron’s outcome selectivity (tuning) was tracked across pairs of consecutive days. From CD to IDS, N=1083 neurons in 17 animals; from IDS to Rev, 1543 neurons in 16 animals; from Rev to EDS, 1382 neurons in 15 animals; from EDS to IDS2, 1443 neurons in 15 animals; from IDS to the first SEDS session, 1519 neurons in 21 animals; from the first to second SEDS sessions, 635 neurons in 21 animals. Neurons included were those with significant modulation by trial outcome (rank sum, p<0.05) in at least one of the two consecutive sessions. Y-axis: Pearson correlation coefficient for outcome selectivity, measured as the difference in mean z-scored activity, averaged over the intertrial interval, between incorrect and correct trials. B. Scatter plot of outcome selectivities for individual, significantly modulated, neurons in the first and second SEDS sessions (same neurons as A). Pearson R=0.49, p=1.6× 10^−39. Correlation for all neurons (unmodulated as well as modulated) was 0.36, p=7.5× 10^−60n N=1959 neurons. C. Similarity of outcome selectivity over multiple SEDS sessions. 173 neurons, from 8 animals, recorded in at least four consecutive behavioral sessions. Neurons significantly modulated by trial outcome in at least one session (rank sum p<0.05) were included, though analysis of all neurons produced comparable results (1091 neurons, R=0.17-0.38, all p values < 1× 10^−7). D. Trial-aligned correlation coefficients comparing outcome selectivity (defined as in A-C) between trial t-1 and trials t-2 through t-6. N=4740 cells from 21 animals. Gray box: stimulus presentation. E. A time-compressed representation of the results from (D), broken out by individual neurons. X-axis: mean activity on incorrect trials; y-axis: mean activity on correct trials. Insets: example cells preferring correct (top) and incorrect (bottom) outcomes in trials t-1 to t-5. F. Schematic diagram of Euclidean distance and vector angle computed from trial-averaged outcome (correct vs incorrect) conditions in neural activity space. G. Left: Euclidean distances for all combinations of trial t-1 and t-2 outcomes in N-dimensional activity space (same data as A-E). Bar heights and error bars are means ± 95% confidence intervals over 500 trial sub-samplings with replacement (50% of trials in one condition, 50% in the other). The rightmost (gray) bar is the mean distance within condition (e.g. [t-1 correct, t-2 correct] vs [t-1 correct, t-2 correct]) across sub-samplings. Right: angles between pairs of trial outcome vectors shown at left. All vectors use [t-1 incorrect, t-2 incorrect] as vertex. Bar heights and error bars are mean ± 95% CI as at left. Leftmost (black) bar is the mean within-condition angle, a proxy for baseline, as more dimensions tend to increase vector angles in noisy data (Brinkman and Charikar, 2005).

We then examined the temporal stability of *outcome* signals over trials. The neurons responsible for coding a given trial outcome might shift between trials, with different groups of neurons inheriting representations of different past trials. Alternatively, these trial histories might be multiplexed by the same neurons, whose activity could be incrementally modulated with each correct and incorrect trial. To test these possibilities, we compared *outcome* selectivities for successive past trials and found that they were indeed supported by correlated populations, whereby, e.g., neurons excited by reward in trial t-1 were also excited by reward in trial t-2 (**Fig. 4D**). As a further illustration of this phenomenon, the path of each neuron’s incorrect/correct ratio over successive past trials tended to display a linear regression toward unity (**Fig. 4E**).

To better understand the evolution of outcome representations over trials, we analyzed the relative locations of these representations in the activity space. The representation of each recent trial might be encoded separately and relayed to downstream brain regions for further processing, or alternatively, they might be compressed into a low-dimensional readout, signaling the animal’s overall performance. An analysis of the Euclidean distances and vector angles of pairs of past outcomes in the neural activity space (**Fig. 4F**) revealed that more similar outcome combinations (e.g. [t-1 incorrect, t-2 correct] vs [t-1 correct, t-2 incorrect]) were less distant than more different outcome combinations (e.g. [t-1 incorrect, t-2 incorrect] vs [t-1 correct, t-2 correct]), and that all pairs of outcome vectors were more colinear (exhibiting lower vector angles) with one another than with vectors from shuffled trials (**Fig. 4G**). Together, these results define a coding scheme in which a subset of PFC neurons represent the outcome of each trial, with their activity state modulated in an incremental, relatively colinear way, with decreasing amplitude as trials recede into the past, rather than different groups of neurons representing outcomes at different latencies from each trial.

### Similar task responsiveness in two major PFC projection neuron populations

Our findings above indicate that PFC cells are highly functionally heterogeneous with respect to their contributions to cognitive flexibility in this set-shifting paradigm. To understand the mechanistic basis of this functional heterogeneity, we questioned whether the task-related activity of a PFC neuron might be a function of its long-range efferent connectivity profile or of its laminar location, a correlate of afferent connectivity. While not mutually exclusive, these two proposed correlates of functional specialization within cortex have motivated numerous studies in recent years, and noteworthy findings have lent support to both hypotheses. Pioneering work by Otis et al, for example, found dissociated response properties in cortico-striatal and cortico-thalamic neurons in a Pavlovian conditioning task, with the former activated and the latter inhibited by reward-predicting conditioned stimuli (Otis et al., 2017). However, because the projection populations examined in that study were located in different cortical layers (layer 2/3-5 and layer 6, respectively), the contribution of laminar depth to coding differences could not be directly addressed. Recent work by Liu et al demonstrated that ventral hippocampal afferents within mouse infralimbic cortex show both projection and layer specificity in their axonal targets, preferentially driving cortico-cortical projection neurons over other projection subtypes, and preferentially driving layer 5 neurons over layer 2/3 neurons (Liu and Carter, 2018). Thus, projection specificity and laminar depth may both contribute to functional specialization of subpopulations within PFC.

To examine the degree to which projection target specificity contributed to the functional heterogeneity of in PFC neurons during set-shifting, we examined the functional properties PFC-VMS and PFC-MDT neurons, two projection-defined PFC output populations whose target structures have both been implicated in supporting cognitive flexibility in prior work (Block et al., 2007; Floresco et al., 2006; Marton et al., 2018). Labeling of both populations within the same animals (Soudais et al., 2001; Tervo et al., 2016), using rAAV-Cre labeling, revealed two largely non-overlapping cell types that were also spatially intermingled (**Fig. 5A**). Despite their being spatially interspersed, double labeling of PFC-VMS and PFC-MDT neurons showed little (<5%) overlap (**Fig. 5B**). Examination of fluorescent axons in fluorescently labeled neurons from the two groups (**Fig. 5C-D**) revealed two distinct populations. PFC-VMS neurons sent dense collateral projections to the claustrum, septo-hippocampal complex, caudoputamen, and basolateral amygdala, but not to MST. PFC-MDT neurons sent dense collateral projections to the habenula, hypothalamus and other thalamic nuclei, but not to VMS **Fig. 5E** and **S13**).

**Figure 5:**
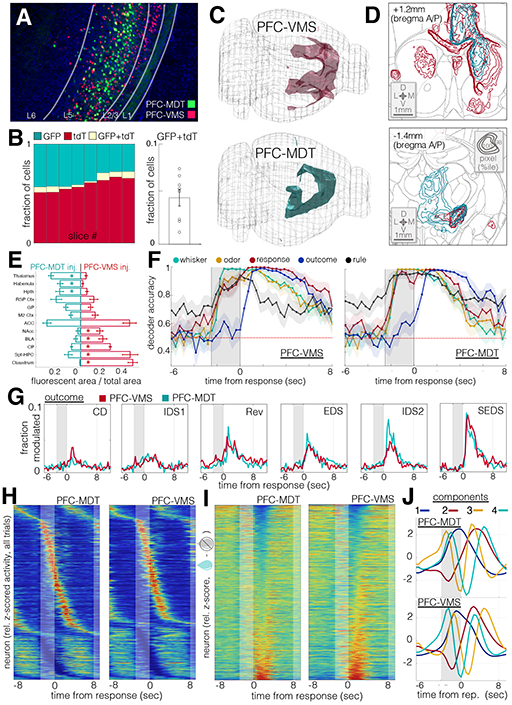
PFC-VMS and PFC-MDT projection populations exhibit similar task response properties. A. Dual PFC-VMS and PFC-MDT projection labeling in a single preparation. Green: rAAV2-CAG-mNeuroGFP in MDT; red: AAV1-hSyn-DIO-nls-tdTomato in PFC and CAV2-Cre in VMS (DAPI in blue). B. Cell counts by slice from the experiment in (A). Left: relative green, red, and double-labeled cell counts. Right: median relative overlap (N = 8 sections, 2 animals). C. Composite images of long-range axon projections from PFC-VMS (N=5 animals) and PFC-MDT (N=5 animals) labeling (see Methods). Volumes shown are thresholded to ≥98^th^ percentile for pixel brightness. D. Coronal sections and contour plots of fluorescent axon density in PFC-VMS (top) and PFC-MDT (bottom) labeled animals. Top: a composite (N=5 animals) of pixel brightness in a coronal cross-section centered on the nucleus accumbens. Concentric contours corresponds with relative pixel brightness (outermost contour, 98^th^ percentile, innermost contour, 100^th^ percentile). Bottom: composite (N=5 animals) of pixel brightness in a coronal cross-section centered on medial thalamus. Contour brightness scale as in (top). E. Mean +/− SEM plots showing the fraction of pixels in each region showing fluorescence above the threshold percentile. 2-way ANOVA revealed an interaction effect between injection type and region (F=17.96, p=0). Post-hoc t-tests revealed significant differences (unpaired t-test p<0.01) for thalamus, habenula, caudoputamen, septo-hippocampal complex, and claustrum, and p<0.02 for hypothalamus, basolateral amygdala, and nucleus accumbens. F. Trial-aligned SVM decoder accuracy for whisker stim, odor stim, response, outcome and rule. N=1115 units from 8 animals (PFC-VMS, left), and N=1770 units from 9 animals (PFC-MDT, right). G. Histograms traces of the fraction of cells modulated (rank sum p<0.01) by trial outcome over trial-aligned timepoints and through learning stages for PFC-VMS (red) and PFC-MDT (green) neurons. H. Trial-averaged activity histograms for PFC-VMS (left) and PFC-MDT (right) cells. Each cell’s mean trace is normalized to its peak value, and cells are sorted by time of peak excitation (top) or inhibition (bottom). I. Mean outcome difference (incorrect – correct) traces for PFC-VMS and PFC-MDT neurons. Each cell’s mean difference trace is normalized to its peak value, and cells are sorted from most strongly preferring correct outcomes (top) to incorrect outcomes (bottom). J. First four principal components for trial-averaged activity histograms in PFC-MDT (top) and PFC-MDT (bottom) neurons.

Contrary to our expectations, the two populations showed a striking degree of overall similarity in their task-responsiveness. We used linear decoders to assess which task-related variables were represented in the population-level activity of the two cell types and found that both exhibited robust representation over similar time-courses for all five examined task features (**Fig. 5F**). Both populations showed modulation by trial outcome that emerged through successive learning stages (**Fig. 5G**). Similar distributions of neurons in each population were excited vs inhibited by trial onset (**Fig. 5H**) and the two cells types each showed comparable distributions of correct-preferring and incorrect-preferring neurons (**Fig. 5I**). Principal component analysis showed similar trial-aligned temporal profiles for the main components (**Fig. 5J**), illustrating similar underlying structures accounting for the largest proportions of variance across the populations. Together, these results show that while individual PFC neurons are highly heterogeneous in their functional properties, efferent projection targets do not account for this functional heterogeneity in these two major projection subtypes.

### Interference with feedback monitoring in PFC projection populations impairs set-shifting

The finding of durable representations of trial feedback signals over successive trials raised the possibility that the task-critical role of the PFC might be feedback monitoring. However, the robust representation of stimuli and task rule, particularly during stimulus presentation, tended to support the prevailing model of PFC involvement in set-shifting, namely that PFC activity controls top-down, attention-mediated biasing of rule-dependent action selection in the task. We next tested these possibilities using projection-targeted optogenetics.

We used the soma-targeted anion-conducting channelrhodopsin stGtACR2 (Mahn et al., 2018) to inhibit PFC activity (**Fig. 6A**). Photoactivation of the channel across a 10-fold range of light intensities produced strong silencing of spiking activity in extracellular recordings (**Fig 6B**). Next, we tested the requirement of PFC activity for successful set-shifting performance in three temporally controlled stimulation regimes: during trials (0.5s trial-ready cue period, 2.5s stimulus presentation, and ≤1.5s response window), during the ITI (8-10s epoch triggered on response lick) following CG trials, or during the ITI following ICG trials (**Fig. 6C**). Trial blocks (beginning with rule switch and ending with the animal reaching criterion for the new rule) alternated light off/on.

**Figure 6:**
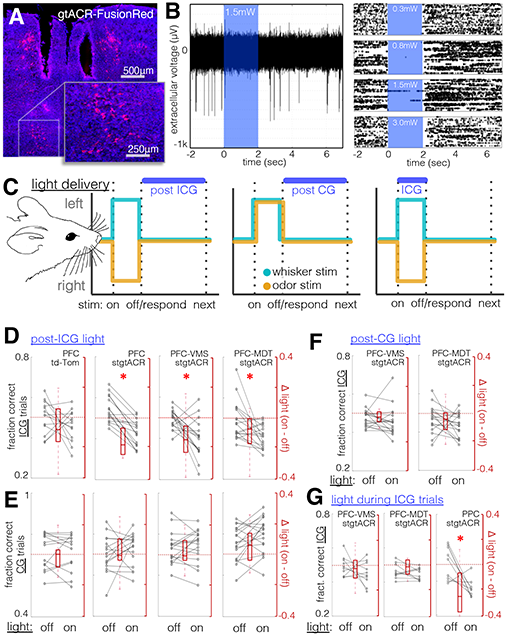
Set-shifting performance requires PFC-VMS and PFC-MDT activity following rule-informative trials. A. Cre-mediated expression of gtACR-FusionRed in PFC-VMS neurons, and fiber tracks from chronically implanted bilateral optical fibers (fixed dissue, DAPI in blue). B. Electrophysiological demonstration of gtACR-mediated silencing. Left: a trace of spontaneous activity (7 seconds), with a 2-second light epoch interposed (1.5mW, 470nm light, 200μm fiber, PFC, see Methods). Right: rasters of 50 sweeps for each of four light intensities: 0.3mW (top), 0.8mW, 1.5mW, and 3.0mW (bottom). C. Schematic diagram of light delivery conditions. Light was delivered during the inter-trial interval following incongruent trials, following congruent trials, or during incongruent trials. D. Effect of light stimulation following incongruent trials on incongruent trial performance (fraction of trials correct). Left: viral control animals (td-Tomato expressed in PFC); second from left: animals with pan-neuronal expression of stgtACR2 in PFC; third from left: animals with stgtACR2 expressed in PFC-VMS projection neurons; right: animals with stgtACR2 expressed in PFC-MDT projection neurons. N = 13, 16, 17, 18 resp. Sign rank p = 0.4, 0.0005, 0.007, 0.01, resp. E. Effect of light stimulation during ITIs following incongruent trials on *congruent* trial performance (proportion of trials correct). Left to right: as in (a) above. N = 13, 16, 17, 18, resp. Sign rank p = 0.9, 0.5, 0.9, 0.07, resp. F. Effect of light stimulation during ITIs following congruent trials on incongruent trial performance. N = 18, 18, resp. Sign rank p = 0.1 and 0.1, resp. G. Effect of light stimulation during trials (stimulus presentation + response window) on incongruent trial performance. N = 10 and 10, resp. Sign rank p = 0.4 and 0.4, resp. Right: Effect of light delivered to posterior parietal cortex (PPC) during incongruent trials on incongruent trial performance (N=9 animals, ranksum p=0.02, opsin-negative control group, N=8 animals, p=0.74).

Photoactivation of pan-neuronally-expressed stGtACR2 during the inter-trial interval following incongruent trials impaired performance on ICG trials (**6D**) but not on CG trials (**Fig. 6E**). No effect of light was seen for control tdTomato-expressing animals (**Fig. 6D-E**). This effect was also seen in animals expressing stGtACR2 in PFC-VMS or PFC-MDT projection neurons (**Fig. 6D-E**). When light was delivered following CG trials, no impairment was seen on ICG trial performance for either projection population (**Fig. 6F**), indicating that the impairment seen with post-ICG inhibition reflected an interference with prior trial feedback, rather than preparation for the subsequent trial. The impairment on ICG trials that resulted from photoactivation of PFC following ICG trials was seen both early and late in trial blocks (**Table S16**). Together, these results confirm a critical role for post-trial activity in both PFC-VMS and PFC-MDT neurons in enabling set-shifting.

Strikingly, neither PFC-VMS nor PFC-MDT activity was necessary for execution of the rule-guided response, as no impairment was seen with photoactivation during ICG trials (**6G**). Failure to disrupt performance by silencing during trials demonstrates that, even though PFC activity represents the task rule and identities of the stimuli (**Fig 2F, 5F**), these representations are not critical for correct responding. In light of this unexpected result, we sought another association area that might mediate rule-dependent responding in real-time. We chose the posterior parietal cortex (PPC), which has been previously implicated in cognitive flexibility (Fox et al., 2003; Prado et al., 2017), as well as in monitoring sensory history (Akrami et al., 2018). Silencing of PPC during trials impaired performance on ICG trials but not on CG trials (**Fig. 6G**), indicating that the PPC mediates responding in the task in a specifically rule-dependent manner, possibly by an attentional mechanism.

### Feedback-related activity follows an anatomical gradient, independent of efferent target

The surprising similarity of both task-related activity and the task-critical function of the PFC-VMS and PFC-MDT pathways left open the question of whether the functional heterogeneity of PFC neurons might be explained by their laminar distribution. We therefore examined whether the relative locations of neurons within the PFC imaging field affected task-related activity. As has been demonstrated in sensory cortex (Smith and Kohn, 2008), temporal correlations across pairs of simultaneously-recorded neurons decayed with distance (**Fig. 7A**), demonstrating an association between spatial proximity and temporal coactivation within the context of the task.

**Figure 7:**
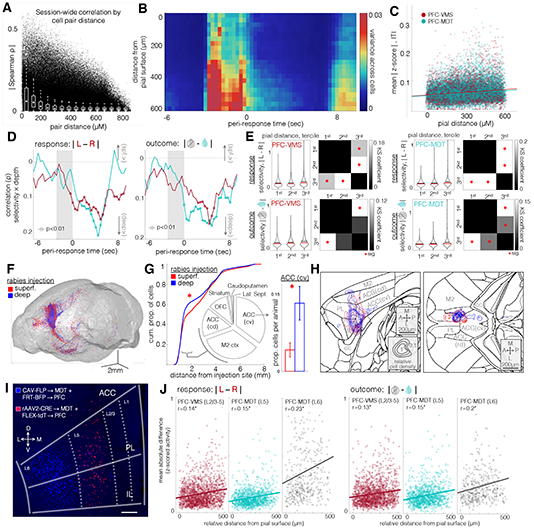
Post-trial feedback-related activity in projection populations is organized by a common topological gradient. A. Spatial distance of all simultaneously recorded cell pairs vs. temporal correlation. Correlations are partial correlations, controlling for changes in putative neuropil (temporal profile background component from CNMF-E source extraction algorithm). Spearman R for distance vs partial correlation is −0.25, p=0 for N = 1562607 pairs from 4740 cells in 21 animals. White plots are box plots for 60μM bins. B. Cross-neuron variance of trial-averaged activity, over trial-aligned timepoints and relative distance from pial surface. Warmer colors correspond with higher variance per spatiotemporal bin. Units are means of z-scored activity. Same cells as in (A). C. Time-averaged data from (B), broken out by neuron and by projection cell type. Correlation of absolute values of z-scored activity with relative pial distance during ITI for PFC-VMS and PFC-MDT neurons. PFC-MDT Spearman ρ = 0.16, p = 8e-79, 1770 neurons, 9 animals; PFC-VMS Spearman ρ = 0.12, p = 3e-52, n=1155 neurons, 8 animals. D. Correlations of feature selectivity with pial distance at trial-aligned timepoints. Left: *response* selectivity; right: *outcome* selectivity. Selectivity is defined as mean absolute difference in z-scored activity between conditions. Circles, Spearman p<0.01. PFC-VMS in red, PFC-MDT in green (same units as in D). Note: y-axis is inverted so that “deep>” values appear on the bottom. E. Distributions of cell feature selectivity for response and outcome during ITI for PFC-VMS and PFC-MDT populations, binned by tercile of relative pial distance. Top row, *response* selectivity; bottom row, *outcome* selectivity; left column, PFC-VMS neurons; right column, PFC-MDT neurons. Red cross-bars are medians, black cross-bars are means. Binned spatial depths were tested for significant differences using a Kolmogorov-Smirnov fit test. For *trial outcome* selectivity across spatial depth, the KS Chi-square for PFC-MDT neurons was 23.09, p=4*10^−05; for PFC-VMS neurons, Chi-square was 19.3, p=0.0002. For *response* selectivity across spatial depth, KS Chi-square for PFC-MDT neurons was 30.2, p=1*10^−06; for PFC-VMS neurons, Chi-square 36.54, p=6*10^−08. Checkered plots are coefficients from post-hoc 2-sample Kolmogorov-Smirnov tests, comparing distribution similarity across all pairs of terciles. Red asterisks are comparisons with Bonferroni-corrected significance. F. Composite brain volume from the rabies tracing experiment. Shown is a three-dimensional rendering of labeled starter cells in PFC, as well as brain-wide retrogradely-labelled afferent cells synapsing onto deep and superficial projection populations (superficial injections, 300μm from midline, 50nL; deep injections, 600μm, 50nL virus). N=45421 cells from 22 animals, 13 deep, 9 superficial injections. Starter cells were PFC-VMS (N=6 injections) and PFC-MDT (N=16 injections). Two way GLM revealed no significant differences in the number of cells labeled according to projection cell type (t=1, p=0.3) or injection depth (t=1.1, p=0.3). G. A Kolmogorov-Smirnov goodness-of-fit test was used to compare distributions of cell density by distance from injection site. Red and blue traces are cumulative density plots for cells labeled by deep (blue) and superficial (red) injections (KS p-value 2.07e-26, n= 9752 and 4755 cells from superficial and deep injections in 9 and 13 animals, respectively). The regions immediately surrounding the injection site (PL, IL, and rostral ACC) were excluded, as they were likely to contain a large number of labeled starter cells. The largest difference in labeled cell density between deep and superficial injections (wherein the first-order derivatives of their CDF curves diverged) was observed in the range of 1mm to 2mm distance from the injection site. Labeled cells in this range were most dense in caudo-ventral ACC (35%), secondary motor cortex (34%), caudo-dorsal ACC (16%), striatum (11%), caudoputamen (2%), and lateral septum (<1%). Of these, the subregion with the most total cells, caudo-ventral ACC, showed a significant difference in cell count by injection depth, with deep-injection animals showing a higher proportion of cells in cv-ACC than superficial-injection animals (0.13 +/− 0.03 vs 0.04 +/− 0.01 of total per animal, t=2.2, p=0.04). H. Cell density contour plots for sagittal (left) and horizontal (right) cross-sections through the regions surrounding ACC. Concentric contours show summed density pooled across all animals (blue, deep-injection animals, red, superficial-injection animals), normalized within the cross-section shown (a.u.). Higher cv-ACC density in deep-injected neurons can be seen in both orientations. Relatively higher density in M2 cortex in superficial-injection animals did not meet significance. I. Labeled subpopulations of PFC-MDT cells with selective tropism for rAAV2 (predominantly located in layer 5, hereafter referred to as PFC(L5)-MDT neurons) and Cav (predominantly located in layer 6, hereafter PFC(L6)-MDT neurons). Dual labeling was achieved by co-injecting a viral mixture of rAAV2-Cre and Cav-FLP in MDT, and a second viral mixture of FLEX-tdT and FRT-BFP in PFC. J. Pial depth and feature selectivity for the three projection subtypes. <u>Right</u>: Kruskal Wallis ANOVA of cell selectivity for *response* across cell types: Chi-square statistic = 440, p=2*10^−96. Post-hoc GLM testing of cell type by pial distance showed significant group differences between PFC-VMS vs PFC(L6)-MDT (t=3.4, p=0.0007), and for PFC(L5)-MDT vs PFC(L6)-MDT (t=3.9, p=0.0001), but not for PFC-VMS vs PFC(L5)-MDT (t=1.0, p=0.3). Post-hoc testing of within-group effect of pial distance on cell selectivity for *response* and trial *outcome* revealed parallel effects (no cell-type-by-pial-distance interaction effects) for all three groups. Spearman R for pial distance by *response* selectivity was significant for PFC-VMS (r=0.14, p=8× 10^−9), PFC(L5)-MDT (r=0.15, p=1*10^−7), and PFC(L6)-MDT (r=0.23, p=1*10^−6). <u>Left</u>: Kruskal Wallis non-parametric anova of cell selectivity for *trial outcome* (mean absolute difference in z-scored activity between correct and incorrect trials during the subsequent inter-trial interval) across cell types: Chi-square statistic = 134, p=10^−30. Post-hoc GLM testing of cell type by pial distance, however, showed no group differences between PFC-VMS vs PFC(L5)-MDT (t=0.5, p=0.6), between PFC-VMS and PFC(L6)-MDT (t=1.25, p=0.2), or between PFC(L5)-MDT and PFC(L6)-MDT (t=1.6, p=0.1). Spearman R for pial distance by *trial outcome* selectivity was also significant for each group: PFC-VMS (r=0.13, p=2*10^−8), PFC(L5)-MDT (r=0.15, p=3*10^−7), and PFC(L6)-MDT (r=0.2, p=4*10^−5). PFC-VMS: N= 1155 cells from 8 animals; PFC(L5)-MDT: N=1770 cells from 9 animals; PFC(L6)-MDT: N=430 cells from 8 animals.

To assess whether neurons’ responsiveness to trials varied as a function of their distance from the pial surface, we quantified the population variance of trial-averaged waveforms to capture the magnitudes of neuron responses, including those both excited and inhibited at each time point. More deeply situated neurons (further from pial surface) showed greater population variance (**Fig. 7B**), exhibiting more diverse trial responses than more superficially located neurons. Greater response *amplitude* (absolute value of z-scored activity) was seen in deeper neurons from both PFC-VMS and PFC-MDT populations **Fig. 7C**).

In addition to trial responsiveness, task-related information was heterogeneously distributed across the cortical laminar axis, with more deeply situated neurons exhibiting greater selectivity for *response* and *outcome* during the ITI (**Fig. 7D)**. General linear models for the neuron selectivities to *response* and *outcome* yielded significant coefficients for neuron depth but not for projection subtype. Because the distribution of these selectivities was positively skewed, we used a nonparametric test to compare sample distributions across groups of neurons from three terciles of pial depth and found that sample distributions differed significantly across depth for both *response* and *outcome* selectivities in both projection types, as evidenced by significantly different Kolmogorov-Smirnov scores between spatial terciles in **Fig. 7E**. We analyzed variance in *response* and *outcome* coding along the dorso-ventral axis, testing for differences between the prelimbic and infralimbic subregions, but we observed no such differences (data not shown). The findings of greater trial responsiveness and greater selectivity for trial *response* and *outcome* in deeper neurons led us to look for a potential mechanism by which deep neurons might exhibit stronger coding of task-critical variables than superficial neurons. To address whether deep and superficial neurons have differential inputs, we employed a rabies tracing approach, targeting EnvA G-deleted rabies-mCherry to PFC projection neurons in spatially restricted deep or superficial injections (**Fig. 7F**). We imaged histological sections from across much of the midbrain and forebrain (bregma AP +3mm to −4mm) and annotated the locations of monosynaptic inputs to the PFC projection neurons. A comparison of the distributions of cells labeled by deep vs superficial injections, as a function of their distance from the PFC injection sites, revealed a significant difference, driven mostly by neurons located between 1mm and 2mm from the PFC injection sites (**Fig. 7G**). An analysis of the regions occupied by these neurons showed them to be most densely located in the caudo-ventral portion of the anterior cingulate cortex (ACC), and a comparison of cell densities from deep and superficial injections in this region showed a higher density of projections to deep PFC projection neurons in this region (**Fig. 7G-H**). Given that ACC has been identified as an area critical for both set-shifting behavior and reward-related feedback monitoring in previous studies (Bissonette et al., 2013; Hyman et al., 2017) this enriched input from caudo-ventral ACC to deep PFC projection neurons provides a potential source of input driving the stronger representation of trial feedback-related information seen in the deeper neurons. Further brain-wide differences in the distributions of inputs to deep and superficial PFC projection neurons can be explored in Supplementary Fig. **S14**.

As a further test of the association between laminar depth and stronger coding of *response* and *outcome* signals, we leveraged the selective viral tropism of two separate PFC-MDT projection populations (**Fig. 7I**), one layer 5 population, for which rAAV2 has strong tropism, heretofore referred to simply as PFC-MDT and hereafter referred to as PFC(L5)-MDT, and one in layer 6, for which canine adenovirus (Cav2) has strong tropism, hereafter referred to as PFC(L6)-MDT (Collins et al., 2018). We used these two viruses as Cre expression vectors to selectively label the two populations with GCaMP6f, and compared their selectivity for *response* and *outcome* during the ITI. If greater distance from the pial surface predicts stronger selectivity for *response* and *outcome*, layer-6 PFC-MDT neurons should show stronger modulation by these two variables compared with layer-5 PFC-MDT neurons or PFC-VMS neurons and should also exhibit a superficial/deep spatial gradient within the neuronal subtype. PFC(L6)-MDT neurons indeed showed greater selectivity for either PFC(5)-MDT or PFC-VMS neurons, as well as exhibiting the same within-population spatial gradient seen in PFC(L5)-MDT and PFC-VMS neurons (**Fig. 7J**). Together, these results reveal a significant source of explanatory variance for the functional heterogeneity seen in PFC neurons during set-shifting. These neurons form a topological gradient, with deeper neurons exhibiting more trial responsiveness and stronger selectivity for trial *response* and *outcome*, potentially driven by differences in their afferent connectivity profiles and in particular by differential innervation by the caudo-ventral ACC.

## Discussion

We began this study with the aim of more clearly resolving the circuit-level mechanisms by which prefrontal activity uses attentional sets to provide context-dependent modulation of sensorimotor processing in set-shifting tasks. The attentional set model of prefrontal involvement in cognitive flexibility, built upon decades of influential literature with thousands of citations, frames the prefrontal cortex as a mediator of top-down cognitive control, which, in the context of an attentional set-shifting task, would facilitate the filtering of multimodal sensory inputs and the biasing of corresponding motor responses in accordance with context-dependent task rules. Instead, what we present is a very different model for PFC involvement in set-shifting. Rather than modulating attention in real-time, PFC output neurons serve to integrate and maintain representations of recent behaviors and their consequences.

While some prior pharmacological silencing studies found PFC activity to be critical for recall but not acquisition of rule switching, and that PFC activity was not critical for rule switches once the rules became familiar (Rich and Shapiro, 2007), those experiments were performed in spatial tasks with changing navigation rules, which may not explicitly engage attention or suppression of irrelevant sensory cues in the same way that a cross-modal set-shifting task does. This difference may account for the result of the muscimol experiment in this study, which *does* find PFC activity to be necessary for switch acquisition, as well as for continued performance of rule shifts after overtraining. We note that the timing of rule shifts, triggered by criterion performance, means that anticipation of these rule shifts cannot be ruled out. However, such a strategy would necessarily involve both acquisition of the new rule as well as anticipation of the change, and it would therefore not eliminate the requirement for any of the key cognitive components of the task. Moreover, the evidence of rule-guided performance (**Figure 1F**) and the failure of animals to immediately abandon the previous rule following a rule shift (**Figure 1G**) argue against such a confound. We also note that, while the training sequence used to expose animals to multiple task rules and stimulus exemplars is based on standard protocols that have been well validated by earlier set-shifting studies (Birrell and Brown, 2000; Bissonette et al., 2008; Tait et al., 2014), we did not specifically test the effect of each training step on the physiology of behavioral strategy used in the final task. Nevertheless, the essential cognitive requirements of the task, as well as the performance data analyzed both with and without perturbation (**Figure 1C-G**), strongly support the conclusion that the protocol achieves comparable results and engages the same cognitive strategy as the protocols on which this task is based.

The finding of retrospective and persistent representation of response and outcome signals, modulated by the demands of the task, are not without precedent. For example, Sul et al found that response on a two-armed bandit task was not represented in PFC activity in advance of the choice (Sul et al., 2010), but this was in an explicit test of reinforcement learning and choice valuation. The attentional set model of PFC involvement in set-shifting predicts that representations of rule, stimuli, and corresponding responses should be activated during decision-making in order to modulate motor responses in downstream structures, such as striatum, thalamus and periaqueduct grey matter.

Responsibility for modulating attention, possibly by active suppression of irrelevant stimuli, may be played by regions with known involvement in multisensory integration – it was this hypothesis that led us to inhibit PPC activity during stimulus presentation. This perturbation did interfere with performance, on ICG but not on CG trials, lending support to the hypothesis.

The finding that *response* and *outcome* are not natively encoded as functions of the overt discrimination behavior, but emerge only once the task rules require repeated/continuous reinforcement learning, lend further support to the conclusion that the role of PFC in this task is to facilitate rule switching by monitoring feedback. These findings build on a body of research showing that feedback signals within PFC are modulated by expectation and by cognitive states, including attention (Smout et al., 2019). Extensive research in feedback-related negativity (Hauser et al., 2013) and reward prediction error (Garrison et al., 2013) support the idea that feedback-related signals in prefrontal areas are dynamically mediated by ongoing task demands and are disrupted in disorders in which set-shifting is also disrupted (Toyomaki et al., 2017).

It is noteworthy that, despite not being critical for task performance, stimulus information is nevertheless robustly expressed in neural activity during stimulus presentation. The observation of strong task-related activity that is nevertheless evidently not critical for task-critical performance, raises the question of what adaptive information-processing purpose this activity may serve, and it underscores the need to assess the function of neural activity through perturbation, rather than assuming a critical role for all task-related signals.

Future work will seek to elucidate potential neuromodulatory and plasticity-related mechanisms behind these signals. Given the high density of input from the cholinergic nuclei of the basal forebrain that synapses on to these projection neurons (**Fig. S14-B**), and the established relationship between the central acetylcholine neuromodulatory system and attention-mediated behavior (Ljubojevic et al., 2014; Proulx et al., 2014), the possibility that acetylcholine plays a key role in mediating these feedback monitoring signals remains strong.

## Acknowledgements

T.S. is supported by NIH (1K99MH117271), the Brain and Behavior Research Institute, and the Leon Levy Foundation. C.L. is supported by NIH (R01 MH109685, MH118451, MH123154) and by the Rita Allen Foundation, Hope for Depression Research Foundation, Pritzker Neuropsychiatric Disorders Research Foundation, and the Foundation for OCD Research. The authors thank S. Fusi, F. Stefanini, and L. Grosenick for consultation on the planning and implementation of data analyses.

## Author Contributions

CL and TS designed the behavioral task and planned the study. TS and MS performed the injection and implantation surgeries and gathered the behavioral data. TS and MS performed the neural recordings. TS analyzed the behavioral and neural data. JK performed the TCA analysis. GMN assisted with the EnvA G-deleted rabies experiment. TS and CL wrote and edited the manuscript.

## STAR METHODS

### LEAD CONTACT AND MATERIALS AVAILABILITY

This study did not generate new unique reagents.

### EXPERIMENTAL MODEL AND SUBJECT DETAILS

Male C57BL/6 mice (Jackson Labs) were used for all experiments, aged 6-10 weeks at first use. Mice were housed in a Weill Cornell Medical College facility and were maintained on a 12-hour light-dark cycle. Except when water-restricted for the purpose of behavioral training and testing, all mice were given *ad libitum* access to food and water. Littermates underwent prism or fiber implant surgeries within the same week, and mice were group-housed with littermates with the same surgery status. All procedures were approved by the Weill Cornell Medicine IACUC. Sample sizes for each experiment were determined using power analysis estimates computed in Matlab, based on anticipated effect sizes that were estimated from previously published reports whenever they were available, and were powered to detect moderate, biologically meaningful effect sizes.

### METHOD DETAILS

#### Surgery

Animals were placed inside a flow box and anesthetized with isoflurane gas (2%) until sedated, at which point they were placed in a stereotax and maintained on 0.5% isoflurane for the duration of the surgery. Scalp hair was trimmed away, and a midline incision was made using fine surgical scissors (Fine Science Tools), exposing the skull. The periosteum was bluntly dissected away and bupivacaine (0.05 mL, 5 mg/kg) was topically applied. For prism implantation, a large rectangular craniotomy was made above left PFC, extending from 1.5mm anterior to 3.7mm anterior, and from 2.0mm lateral (left) to 0.2mm lateral (right, across midline).

A 0.5-mm burr (Fine Science Tools) and a high-speed hand dental drill (Osada) were used, taking great care not to compress brain tissue or damage the sagittal venous sinus. In the event of venous bleeding, Gelfoam (Pfizer) was applied to the dura surface to accelerate clotting. Gentle irrigation with phosphate-buffered saline (137 mM NaCl, 27 mM KCl, 10 mM phosphate buffer, VWR) was used to clear debris at regular intervals. The dura beneath the craniotomy was removed using the tip of a 26g insulin syringe (VWR) and fine forceps (Fine Science Tools).

Chronically implanted microprisms (1.5mm × 1.5mm × 3mm; M/L,A/P,D/V), from OptoSigma (BK7 borosilicate glass with aluminum hypotenuse and silicon dioxide coating), were implanted at a depth of 2.3mm ventral to brain surface using a stereotaxic micromanipulator (Kopf). During implantation, the prism was held in place using vacuum suction via an 18G blunt needle. As in previous studies (Andermann et al., 2013; Low et al., 2014) minimal reactive gliosis was seen in the coronal imaging field, and maximum calcium-mediated fluorescence was seen 50-150μm past the prism face; therefore imaging planes were confined to this depth.

For PFC-VMS and PFC-MDT projection targeting, an additional craniotomy was made at 1.25mm/1.25mm A/L or 1.2mm/0.35mm P/L, respectively. All head-fixed animals received custom-machined stainless steel head plates affixed to skull surface with Metabond dental cement. Head plates featured a circular central aperture centered around the imaging field (9mm I.D.), with right and left securing arms (25mm total width) that accommodated 0-80 socket screws (0.38g in total). Sterile eye lubricant (Puralube, FischerSci) was administered to prevent corneal drying, and a microwavable heating pad (Snugglesafe) was used to maintain body temperature. Metacam (1 mg/kg, i.p.) was administered after surgery as a prophylactic analgesic.

#### Viral Transduction

AAV of titer exceeding 10^12^vg/ml (Vector Biolabs, UNC Vector Core and Addgene) was used to package the plasmids. For imaging experiments, AAV1-hSyn-GCaMP6f or AAV1-hSyn-DIO-GCaMP6f (Chen et al., 2013) was targeted to PFC. Injection coordinates for PFC were 1.75mm anterior. Two parallel injection tracks were made at 0.2mm and 0.5mm lateral, and in each of these tracks, two D/V sites received infusions, at 2.0mm and 1.5mm ventral to brain surface. For rAAV2-Cre projection targeting, VMS injections were delivered at 1.25mm/1.25mm/4.7mm A/L/V, and MDT injections at 1.2mm/0.35mm/3.2mm P/L/V. Hamilton syringes and beveled 36G or 33G NanoFil needles (WPI) were used, and at each site, the needle was allowed to sit 5min to allow for tissue settling before infusion. Virus was infused at a rate of 50nL/min, for a total of 250nL per site.

For the dual labeling tracing experiment (Fig. 5B), rAAV2-mNeuroGFP was injected in MDT (500nL), rAAV2-Cre was injected in VMS (500nL), and AAV1-NLS-tdTomato was injected in PFC (500nL).

#### Muscimol silencing

Animals received chronically implanted 26G bilateral stainless steel guide cannulae (Plastics One), implanted at 1.75mm/0.35mm/1mm A/L/V. After undergoing the training and task transition sequence up to and including Reversal (see behavioral training protocol below), animals underwent muscimol infusion.

In a familiar cage, bilateral internal infusion cannulae were inserted into the guide cannulae, protruding 0.5mm from the end of the guide cannulae, and were left for 5min to allow tissue to settle. Muscimol (1μg/μL) or physiological saline (0.9%) was infused at a rate of 50nL/min, for a total of 0.25μL. Five minutes after completion of the infusion, the internal cannulae were removed and the mouse immediately began behavioral testing.

#### Optogenetic implantation and stimulation

Bilateral fibers were implanted over PFC (Thorlabs dual fiber cannulae, 700μm center-to-center spacing, 200μm core) at 0.35mm lateral, 1.75 anterior, 1.2mm ventral to brain surface. In the posterior parietal cortex inhibition cohort, in order to cover the medial-lateral extent of PPC, a dual-fiber cannula (700μm center-to-center spacing) was placed over each hemisphere, so that both right and left PPC each had two 200μm-core fibers. The dual fiber cannulae were positioned on the brain surface with the dura intact, at a ML angle of 10° away from midline, with the medial fiber at AP −1.8mm, ML +1.0mm, and the lateral fiber at AP −1.8mm, ML +1.7mm. Light was delivered via Thorlabs M470F3 fiber-coupled LED, 1.5-3mW light power from each fiber. In the trial-concurrent stimulation condition, photoactivation was initiated at the same time as the 500ms trial-ready cue and persisted through the 2.5s stimulus presentation epoch until either a lick terminated the trial or the 1.5s response period expired. In the inter-trial interval stimulation conditions, photoactivation was initiated concurrently with the end of the trial (first lick within the response window or upon the conclusion of the 1.5s response window without a lick) and persisted through the 8-10s post-trial epoch, terminating with the onset of the subsequent trial-ready cue. In the experiment testing the effects of inhibition during the beginning of the inter-trial interval, the photoactivation was initiated concurrently with lickspout selection and persisted for 4.5 seconds before terminating; in the experiment testing the effects of inhibition during the end of the inter-trial interval, photoactivation was initiated beginning at 4.5 seconds following lickspout selection, persisting for the remainder of the 8-10s inter-trial interval. Light was delivered in alternating trial blocks, beginning with the second block of the session (no light delivered in the initial block). Sessions in which animals failed to reach criterion on the initial block were re-run until the animal reached criterion.

#### 2-photon imaging

2-photon calcium imaging (Denk et al., 1990; Nakai et al., 2001) was performed via an Olympus 10x 0.6NA objective, with 8mm working distance. All images were acquired using a commercial two-photon laser-scanning microscope (Olympus RS) equipped with a scanning galvanometer and a Spectra-Physics Mai Tai DeepSee laser tuned to 920nm, operating at 300-500mW. Fluorescence was recorded through gallium arsenide phosphide (GaAsP) detectors using the Fluoview acquisition software (Olympus) using a green light emission bandpass filter (Semrock). Imaging sessions began by performing an isosbestic anatomical scan (810nm 2P excitation light) to aid in relocating the same sites over multiple sessions. Calcium signals were acquired at 256 × 130 pixel resolution, covering a 1500μm × 760μm field of view, with a μm/pixel ratio of 5.85. The scan time was 346ms, with a frame rate of 2.89Hz. All calcium imaging experiments occurred in awake mice. For analysis of SEDS sessions, sessions were concatenated using CellReg (Sheintuch et al., 2017) and non-rigid spatial transformation. Neurons were modeled with a maximal centroid distance of 15μm and a threshold correlation of 0.65. Z-scoring of deconvolved activity traces was performed prior to concatenating across sessions. SEDS neural data sets for each of 21 animals are comprised of a median of 3 sessions (2.5,4 upper, lower quartiles), 1462 trials (1034, 1667 upper, lower quartiles), and 11 set shifts (8.5, 17.5 upper, lower quartiles).

#### Image processing

Videos were motion-corrected using NoRMCorre (Pnevmatikakis and Giovannucci, 2017) implemented in Matlab (Mathworks), and a constrained non-negative matrix factorization-based source extraction method was used to denoise, deconvolve and demix the videos to extract neural traces (using the extensively validated CNMF-E package (Pnevmatikakis et al., 2016) with OASIS signal deconvolution (Friedrich et al., 2017). Sources were well separated from both neighboring sources and surrounding neuropil, as assessed by performing PCA analysis of putative source and surrounding pixels over time and quantifying their isolation distances (Schmitzer-Torbert et al., 2005; Stringer and Pachitariu, 2019). The resulting traces were deconvolved calcium traces corresponding with estimated event rates, which were then z-scored over the full session to normalize. Calcium signals from sequential sessions were concatenated using non-rigid co-registration of spatial cell footprints using CellReg (Crowe and Ellis-Davies, 2014; Sheintuch et al., 2017), (see **Fig. S3–S10** for example traces and quality control metrics).

### Behavioral Training and Testing Protocols

Animals recovered 14 days from surgery before being placed on water restriction for four days, after which they typically drank 1-1.2mL/day. Behavioral training and testing procedures were carried out 5 days per week, with each animal undergoing one session per day. Animals underwent two days of hand-feeding, in which they were handled for up to ten minutes by the experimenter while receiving water from a 1mL syringe with a rounded stainless steel gavage needle.

#### Habituation-1

Following two days of hand-feeding, animals underwent a *Habituation-1* session in the behavioral apparatus, which consisted of an aluminum restraint tube with dual lickspouts positioned at one end. The restraint tube was 27mm in diameter, a width calibrated to allow the animal to groom and adjust its posture during sessions but which prevented significant lateral or vertical body movement. During *Habituation-1*, lickspouts were alternately armed so that a single lick would trigger delivery of a 3uL water bolus, and the identity of the armed lickspout changed every 1-4 trials, with a 1.5s timeout after each bolus delivery. This alternating schedule forced the animal to explore both lickspouts equally to maximize rewards. Animals would periodically venture out of the restraint tube to explore the behavior chamber, at which point the experimenter would guide them back into the tube by hand. The session terminated when the animal stopped licking for ~2min, and the animal was considered to have passed this stage when it had consumed 500μL in a session.

#### Habituation-2

After passing Habituation-1, animals underwent a *Habituation-2* stage, which consisted of the same lick/dispense schedule as Habituation-1, but during which the animal was head-restrained for the first time. Here again, the animal needed to consume 500μL to pass.

#### Habituation-3

After Habituation-2, animals underwent *Habituation-3*, in which lickspouts alternated every 20 lick/delivery trials, to establish left/right trial blocks. As with Habituation-1 and −2, animals were considered to have passed this stage after consuming 500μL of water.

#### Shaping-1

After this, animals underwent a *Shaping-1* session, which introduced the trial structure to be used for the rest of the sessions: a 500ms white noise trial-start cue was presented, followed by a 2.5s stimulus (whisker or odor, depending on the training sequence, which was counter-balanced across animals in the calcium imaging experiment), followed by a ≤1.5s response window, during which the correct lickspout was armed. When the response rate fell to between 0.3 and 0.6 within a 10-trial moving window, reward would be dispensed regardless of whether the animal licked, in order to promote licking. Trials were blocked (20 right, 20 left), and a lick to the incorrect side would not terminate the trial; the animal was allowed to continue until it licked the correct spout. At the start of each trial block, a single 3uL reward was dispensed at the now-rewarded spout. Criterion for passing *Shaping-1* was 500μL consumed.

#### Shaping-2

*Shaping-2* differed from *Shaping-1* in that trials were randomized between sides rather than blocked. Criterion here, and for all remaining sessions up to SEDS, was reaching 80% correct within a 30-trial moving window, and simultaneously performing above 50% on both left and right trials, at any point within the session.

#### Direction bias correction

To counteract animals’ tendency to sporadically exhibit stereotyped direction bias, left/right stimulus probability was dynamically computed as a function of the animal’s concurrent direction bias coefficient. A 10-trial moving window was populated with a [-1] for each incorrect left response, a [1] for each incorrect right response, and a [0] for each correct response. For each trial, the coefficient was the mean over the 10 previous trials. A coefficient absolute value between 0 and 0.25 resulted in 50% left-right stimulus probability; an absolute value between 0.25 and 0.75 resulted in a 33% chance of a stimulus toward the direction of bias; an absolute value between 0.75 and 1 resulted in a 17% chance of a stimulus toward the direction of bias. Trial runs with lick bias coefficients greater than 0.75 were therefore excluded from analysis.

#### Training on discrimination and cognitive shifting tasks

##### Stimuli

Of the 21 mice included in the imaging experiment, 14 underwent initial training with whisker stimuli, and 7 underwent initial training with odor stimuli. Subsequent comparison of stimulus encoding did not indicate any discernible differences between these two groups following the extradimensional set-shift. It should be noted that the hearing range of mice is 1-80kHz (Turner et al., 2005), which falls outside the range of the stimuli used to vibrate the whiskers.

For animals undergoing SD with whisker stimuli, stimuli consisted of a 35Hz sinusoidal stimulus and a 155Hz sinusoidal stimulus. The 35Hz and 155Hz stimuli were delivered bilaterally. These stimuli were generated by a pair of miniature base-frequency audio speakers (2" diameter) coupled to a pair of plastic funnels which served to condense the sound waves into a 5mm diameter compression wave (75 dB). These stimuli produced oscillatory deflections of a majority of whiskers on the order of 10°, and active whisking was routinely observed during these epochs.

Odor stimuli consisted of murine-appetitive oil extracts. For the 7 mice undergoing initial training with odor stimuli, these stimuli were olive oil and sesame oil. The odor port was positioned 5mm below the animal’s nostrils, with the airflow directed up toward the nostrils. Air was delivered through this port at a rate of 2.5 liters per minute through 1/32" inner-diameter polytetrafluoroethylene tubing. Outside of the odor stimulus epochs, clean air was delivered continuously through the odor port. At the onset of the odor stimulus epoch, air was rerouted through chambers containing the oil extracts, so that clean air was completely displaced by air from the oil container within 30ms.

Each trial began with a 500ms audible white noise (2kHz-17kHz) to indicate the start of a trial. Immediately following this cue, a whisker and/or odor stimulus (depending on the task phase as outlined in the table in Figure 1 and descriptions below) were presented for 2.5sec. The animal was permitted to lick the water spouts freely during stimulus presentation, though these anticipatory licks did not terminate the stimulus or trigger reward. Following the 2.5sec stimulus delivery, there was a response window of up to 1.5sec during which a lick to either port would end the trial, trigger delivery of a 3μL water droplet (on correct trials), and begin the inter-trial interval. As shown in Figure 1B, licking toward the ultimately chosen lick port began well before the termination of the stimulus, and 95% of responses came within the first 0.346ms of the response window, a latency that corresponded with the first frame of imaging following stimulus termination.

##### Simple Discrimination (SD)

For the 14 mice undergoing initial training with whisker stimuli, on each trial a bilateral whisker vibration stimulus (one of two possible patterns) was delivered. Through water reward-mediated reinforcement, mice learned to lick the left lick port in response to the 35Hz stimulus and the right lick port in response to the 155Hz stimulus. For the 7 mice undergoing initial training with odor stimuli, animals learned to lick left in response to an olive oil odorant and right in response to a sesame oil odorant. Sessions consisted of up to 350 trials, and sessions were terminated early if an animal failed to respond to ten consecutive trials. An animal was considered to have reached criterion for the SD phase if at any point during a session it reached 80% correct performance, and simultaneously ≥50% correct on both left and right trials, within a 30-trial moving window. The SD phase tests the animal’s ability to learn a stimulus-reward contingency and discriminate between two stimuli from the same dimension (i.e. two odors or two whisker stimuli). When criterion was reached, the current session continued to its end (up to 350 trials), and the animal was moved on to the subsequent task phase on the following session.

##### Compound Discrimination (CD)

An irrelevant stimulus (odor stimulus for mice initially trained on whisker stimuli and vice versa) was added on each trial. Both relevant and irrelevant stimuli were randomly and independently chosen for each trial. The same criterion standard was used as with SD: 80% correct performance (and at least 50% on both right and left trials) within any 30-trial moving window.

##### Intradimensional Shift (IDS)

A new pair of relevant stimuli replaced the stimuli used for SD and CD, while the irrelevant stimuli remained unchanged. For animals initially trained on whisker stimuli, the new stimuli were a 210Hz sinusoid (lick-left) and a train of Poisson-distributed square-wave clicks averaging 210Hz (lick-right). For animals initially trained on the odor rule, the new stimuli were almond oil (lick-left) and orange extract (lick-right). This phase tests the animal’s ability to learn a new stimulus-reward contingency and discriminate between two new stimuli from the same task-relevant dimension. The criterion for passing IDS was the same as for SD and CD.

##### Reversal (Rev)

The same set of relevant and irrelevant stimuli were used as in IDS, but the left-right mapping was reversed. This phase requires the animal to suppress and replace a previously learned stimulus-response contingency. Criterion was the same as in SD, CD, and IDS.

##### Extradimensional Shift (EDS)

The same whisker and odor stimuli were used as in IDS and Rev, but the previously irrelevant modality now became the relevant modality and vice versa. Because relevant and irrelevant stimuli were randomly and independently chosen on each trial, 50% of trials were congruent and 50% were incongruent. This phase requires the animal to associate a new stimulus *category*, the previously irrelevant stimulus modality, with a left-right response mapping. Criterion was the same as in all preceding task phases.

##### Intradimensional Shift 2 (IDS2)

The stimuli in the newly relevant modality were replaced, while those in the newly irrelevant modality were unchanged. This phase serves to ensure that the animal has learned to associate left-right stimulus mappings with multiple exemplar sets in each modality, thereby establishing an abstract attentional set within the newly relevant modality. Criterion was the same as in all preceding task phases.

##### Serial Extradimensional Set-Shifting (SEDS)

For each SEDS session, the relevant stimulus modality was chosen randomly at the start of the session. Stimuli during these sessions were the same stimuli used in the IDS2 phase (whisker: 210Hz sinusoid, left, and Poisson click train, right; odor: almond oil, left, and orange extract, right). Upon reaching criterion (still 80% correct and 50% on both left and right in a 30-trial moving window), the modality rule was automatically switched. This phase tests the animal’s ability to integrate trial feedback to flexibly switch between task rules. As with previous phases, relevant and irrelevant stimuli were randomly and independently generated on each trial. To maximize the number of set-shifts in each session, sessions continued until animals reached satiety (ten consecutive non-response trials) or consumed 1.1mL of water.

### Fixed tissue processing and imaging

Animals were transcardially perfused with 4% paraformaldehyde and 1x PBS. Heads were removed and incubated with all tissues intact in 4% PFA at 6℃. Brains were dissected after 24 hours and fixed in 4% PFA for an additional 24 hours before being dehydrated in a 30% sucrose solution (24-72 hours, until submerged). Tissue was sectioned coronally (in 45μm sections) at −20℃ and submerged in PBS until mounting. (When not in the cryostat tissue was stored at 6℃). Samples were mounted on 25× 75×1 mm slides (Cole Parmer or Fisher Superfrost) with Sigma brand, DAPI-infused Fluoroshield mounting medium. For the L5 vs L6 PFC-MDT dual labeling experiment, Fluoroshield without DAPI was used, to enable BFP visualization.

Imaging was carried out on the Leica DM 5500 B microscope with a Leica DFC360 FX fluorescent camera (7x magnification) and EL600 light source, using a 10x apochromat objective with an NA of 0.4. Images were taken in two channels - the red channel used a CY3 filter cube (excitation filter 545/40, dichroic mirror 565, emission filter 610/76) at a 90ms exposure. The blue channel was captured with a DAPI filter cube (excitation 360/40, dichroic 400, emission 470/40) at a 100ms exposure. Both channels were imaged with full laser power and no gain. To allow for comparison across brain regions, both within and across animals, these imaging parameters were held constant for images gathered for the rabies tracing and fluorescent axon tracing experiments.

Images were acquired using the Leica Acquisition System (LASX) (Version 3.6) navigator feature to capture each sample as a series of 1281.71 × 957.36 um (1392× 1040 px) tiles, which were stitched (with no overlap or blending) by the software during acquisition to produce the complete image.

#### Rabies tracing

Monosynaptic inputs to PFC-VMS and PFC-MDT projection neurons were labeled by using a cross-sectional three-virus approach. At 4 weeks prior to sacking, AAV encoding the TVA rabies B19 glycoprotein (pAAV-EF1a-FLEX-TVA/B19) was injected into PFC. Virus was infused using a 36g beveled NanoFil needle (WPI) and a 10μL Hamilton syringe. Infusions of 250nL were made at each of four sites within PFC: AP +1.75mm, ML −0.2mm and −0.5mm, DV −2.0mm and −1.5mm (brain surface). Virus was infused at a rate of 50nL/min. In the same surgery, retrogradely transported rAAV2-CAG-Cre was injected into MDT (AP − 1.2mm, ML −0.35mm, DV −3.2mm) or VMS (AP +1.25mm, ML −1.25mm, DV −4.7mm) in a volume of 500nL.

Following the dual infusions, Vetbond tissue adhesive was applied to the craniotomy over the exposed dura, and the skin was sutured into place using 6-0 silk sutures. Three weeks later, a second infusion surgery was performed to infuse EnvA G-Deleted Rabies-mCherry in PFC. Using a Nanoject nanoliter injector and a borosilicate pulled glass micropipette, 50nL of virus was infused at either a cortically superficial (ML −0.3mm) or deep (ML −0.6mm) site (AP +1.75, DV −1.7). The animal was again sutured and allowed to recover 1 week before sacking.

#### Fluorescent axon tracing

Surgical preparation, craniotomy, and infusion procedures were the same as those for GCaMP and stGtACR2 projection labeling experiments. AAV1-CAG-DIO-tdTomato was infused in PFC at four sites: AP +1.75mm, ML −0.2mm and −0.5mm, DV −2.0mm and −1.5mm, at a volume of 250nL per site and a rate of 50nL/min. rAAV2-CAG-Cre (500nL) was infused in MDT (AP −1.2mm, ML −0.35mm, DV −3.2mm) or VMS (AP +1.25mm, ML −1.25mm, DV −4.7mm) in a volume of 500nL. Animals were sutured and allowed to recover for four weeks before sacking.

#### Image processing of rabies-labeled and fluorescent axon histology

Coronal slices were mounted and imaged from ~AP +3mm to ~AP −4mm, spanning the frontal pole of the neocortex to the ventral/caudal hippocampus and surrounding regions. Imaging was performed at 1μm/pixel resolution and tiled using Leica software. Image cropping, alignment, de-warping, registration to the Allen Common Coordinate Framework (Wang et al., 2020), and annotation of cell locations was performed using custom Matlab algorithms adapted in part from the cortex-lab/allenCCF package (Shamash et al., 2018). Histological images were mapped to corresponding atlas sections by annotating images with manual control points and generating piecewise regression image transformations (Matlab *fitgeotrans*). Individual labeled neurons were then individually annotated in Matlab and mapped onto atlas locations using the Allen CCF structure tree (data.cortexlab.net/allenCCF).

### Electrophysiological recordings

#### stGtACR2 Photoactivation

AAV1-hSyn-stGtACR2-FusionRed was infused in PFC (AP +1.75mm, ML −0.2mm / −0.5mm, DV −2.0mm / −1.5mm, 250nL/site). Animals were sutured and allowed to recover for 4 weeks. On the day of the recording session, they were anesthetized under isoflurane (2%) and transitioned to 0.5% isoflurane over a 20min monitoring period. The craniotomy from the earlier viral transfusion was widened to an area of ~2mm × 2mm. A 32-channel silicon probe, coupled to a 200μm-core, 0.6NA optical fiber (H6b-style optrode, Cambridge Neurotech) was lowered into the brain at a rate of ~100μm/min to a depth of 1.7mm ventral at the tip. Recording were gathered using an Intan analog-to-digital acquisition system. Spikes were sampled at 20KHz with a 1Hz high-pass filter and 60Hz notch filter, and recording channels were referenced against an alligator clip secured to the neck muscle. Raw voltage recordings were later bandpass filtered between 250Hz and 20KHz using Matlab’d *bandpass* function, and multiunit spiking activity was thresholded at − 0.1mV.

#### Measuring muscimol-induced silencing

Anesthesia was administered as described above in the photoactivation experiment. A larger (~2mm ML × ~3mm AP) craniotomy was opened over frontal cortex. The animal’s head was raised in order to provide a flat (rather than sloped) cortical surface over frontal areas. A multielectrode array (Innovative Neurophysiology Inc.) was arrayed in two rows of 8, spaced at 150μm between columns and rows. The rows were arranged along the AP axis, with the most anterior recording site at AP +2.0mm, DV −0.5mm. and the two rows at 0.3mm and 0.5mm left of the midline. Muscimol (1μg/μL) was infused at a rate of 50nL/min, for a total of 0.25μL, through a 36g needle positioned in between the most anterior pair of electrodes and angled 10° toward the anterior. Spontaneous activity during and following infusion was recorded and processed with the same settings as in the photoactivation experiment described above.

### STATISTICAL ANALYSIS

#### Tensor component analysis (TCA)

To estimate latent population activity associated with each task variable on a trial-to-trial basis, we used an unsupervised tensor component analysis (TCA) approach (Williams et al., 2018) To avoid artifactual sources of variance introduced by session registration, this analysis was performed independently for each mouse and session. Trial-aligned z-scored activity traces (trial × neuron × time) were first normalized to a nonnegative range and smoothed with a gaussian kernel. They were then decomposed into a set of related trial, neuron, and temporal factors. We evaluated models using both standard TCA and nonnegative TCA, in which weights are constrained to positive values, ultimately selecting nonnegative TCA as it provided a more consistent fit. The trial components of these factors were then regressed onto the five primary task variables (whisker, odor, response, outcome, and rule) using Matlab’s glmfit function. Factors that were significantly modulated (Bonferroni corrected t-test) by a given variable in the trial domain were then pooled to estimate temporal profiles for latent population activity underlying each task variable. Tensor reconstruction was performed by performing matrix multiplication on the component neuron*factor, trial*factor, and time*factor matrices. Temporal components shown in Figure 2G are from factors found to be associated with each task variable with a p-value < 10e^10^ using the GLM. This stringent criterion was used in order to include only the top 5% of most reliably modulated factors. Less stringent criteria resulted in the inclusion of factors with a high variance among temporal components.

#### Linear decoding with SVM maximum margin classifiers

Population decoding was performed with maximum-margin linear decoders (Matlab fitcsvm). At each iteration data were separated into equally sized training and testing sets, and in each set, trials were separated into equal numbers from each binary value for the feature of interest. To combine inputs from neurons across multiple sessions, and to remove inter-trial variability each of these trial groups was than randomly separated into 16 trial groups, each of which was averaged into one *super-trial* to serve as a decoder input (16 super-trials for one value of the feature of interest and 16 from the opposing value). This number was calibrated to ensure robust trial subsampling within training and testing sets while removing sufficient trial variability to permit accurate decoding. Standard deviations for classification accuracy were obtained by iterating randomized training and testing over 500 subsamplings with replacement. Decoder testing performed on data from inter-trial intervals was run on binned activity values averaged over the 8 second post response epoch unless otherwise specified.

#### Principal components analysis (PCA)

PCA was used to identify the trial-aligned activity patterns responsible for the greatest variance within the activity space of the neural traces (Fig. 5J). PCA was performed using Matlab’s native *pca* function, and the first three resulting components were plotted through trial-aligned time. PCA was performed separately on trial-averaged traces from PFC-VMS and PFC-MDT sets (neuron × time).

#### General Linear Model

Matlab’s *fitglm* function was used to fit a GLM model to projection cell type and topological location (depth from pial surface). Fitglm was also used to model multiple single neuron activity as a function of 7 task variables: whisker stimulus, odor stimulus, response direction (current trial), response direction (previous trial), trial outcome (current trial), trial outcome (previous trial), and task rule. Unless otherwise specified, GLMs were modeled with normal distributions, as linear functions (no interaction effects) with y-intercept terms, and with identity linkages. For analysis of individual neuron responses to task variables, a separate GLM was computed performed for each time point in the trial independently.

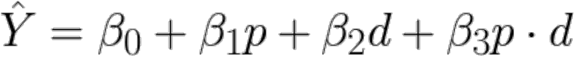

#### Euclidean distance and vector angles in neural activity space

Euclidean distance between pairs of trial averaged conditions was computed as the square root of the sum of the differences between the two conditions across all neurons in the set.

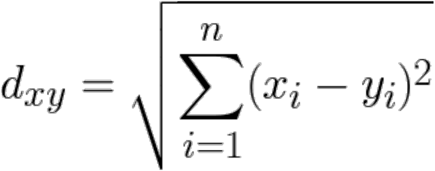

Euclidean distance was also computed on activity vectors scaled by the vector weights of their corresponding SVM decoders in order to scale the amplitude of feature-selective cells relative to noise. Vector angles were computed as the dot product of the difference vectors, divided by the product of their norms. This value was converted into an angle of degrees by taking its inverse cosine.

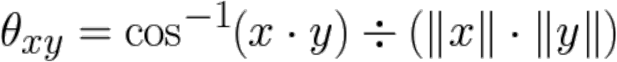

### DATA AVAILABITY

All data and Matlab and Python analysis scripts are available from the corresponding author upon reasonable request.

## Supplemental Figure Titles and Legends

**Figure S1.**
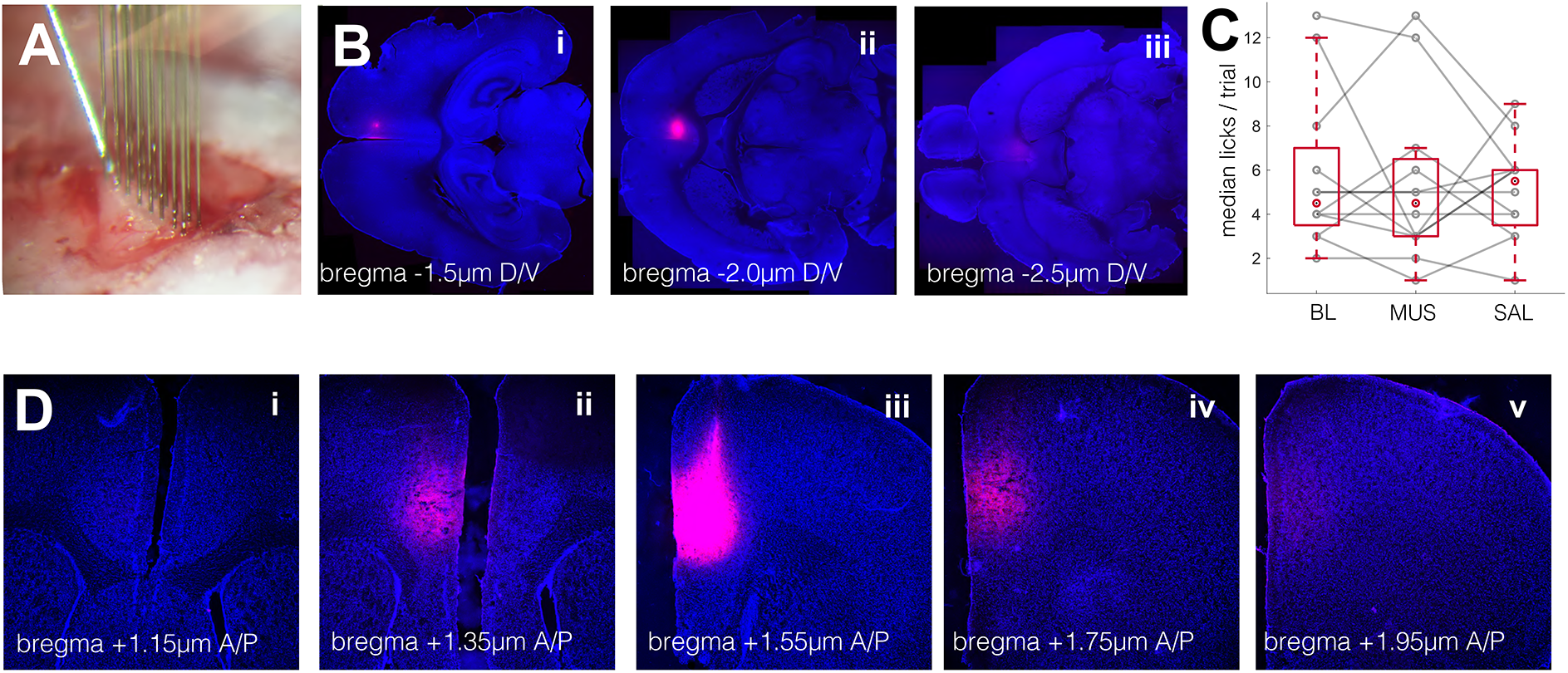
Muscimol infusion experiment. A. 16-channel multi-electrode array (Innovative Neurophysiology Inc.) used to measure the spatial extent of muscimol silencing effects. Photograph taken through a binocular surgical stereoscope (Kopf). The diagonal needle in the upper left is the infusion needle (36g blunt, NanoFil), tilted to an anterior angle of 10° off vertical. The needle was positioned between the anterior-most pair of electrodes. Electrode rows were spaced at 150*μ*m apart, 150*μ*m spacing within rows. B. Histological sections from a fluorescent muscimol injection at three horizontal planes. Red: Muscimol-TMR-X (Sigma Aldrich); Blue: DAPI. C. Comparison of overt licking behavior in animals receiving muscimol, saline, and no injection. Rank sum for the BL-MUS comparison: p=0.88. D. Histological comparison from a fluorescent muscimol injection at five coronal planes. Red: Muscimol-TMR-X (Sigma Aldrich); Blue: DAPI.

**Figure S2.**
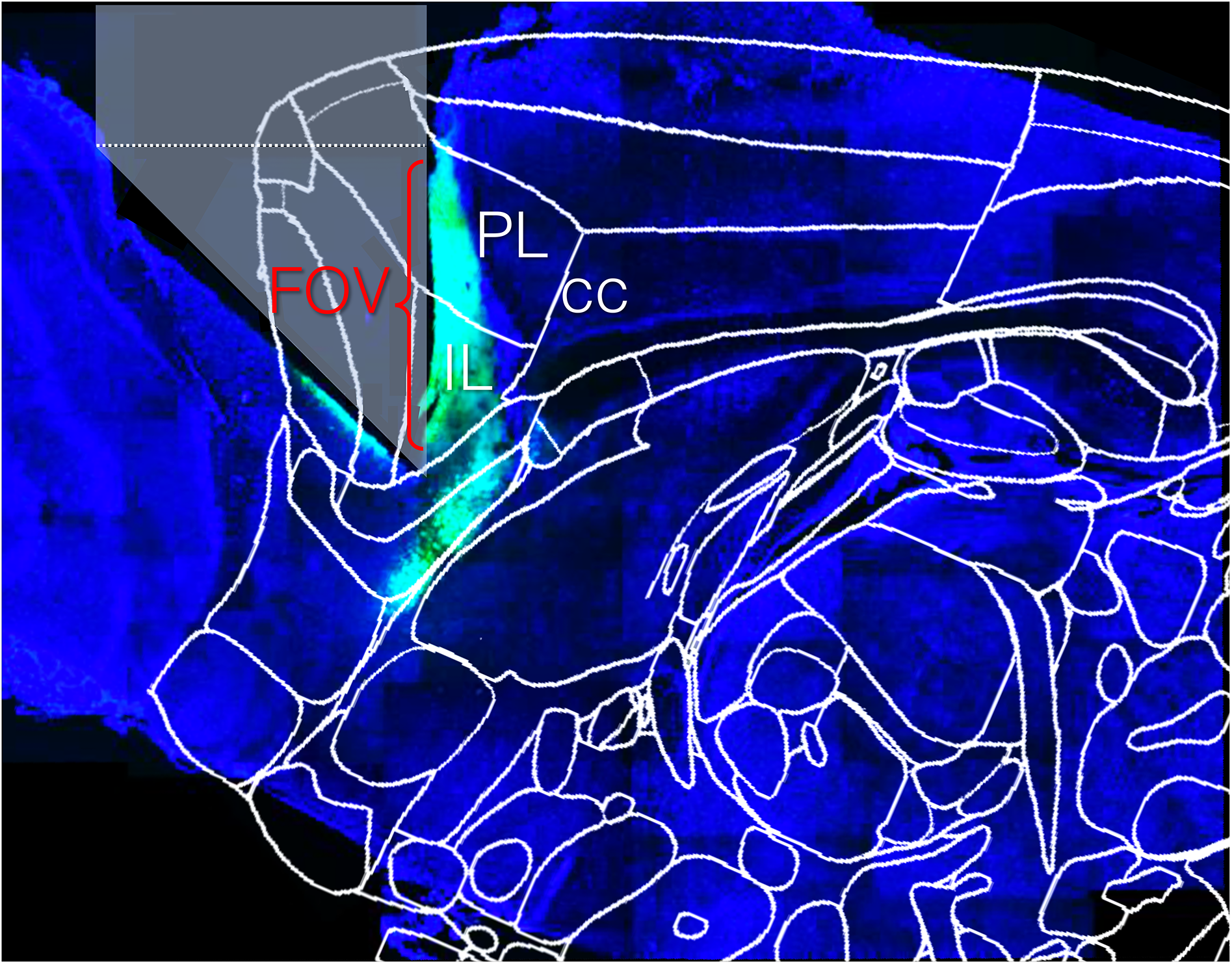
Sagittal section from prism-implant. hSyn-GCaMP6f or hSyn-DIO-GCaMP6f was injected at A/P +1.75mm, M/L −0.2mm and −0.5mm, D/V − 2.0mm and −1.5mm. An atlas image from M/L −0.42mm was superimposed, with the corpus callosum (cc) used to co-register the atlas with the histological image. PL: prelimbic cortex; IL: infralimbic cortex; ACC(rd): rostro-dorsal anterior cingulate

**Figure S3.**
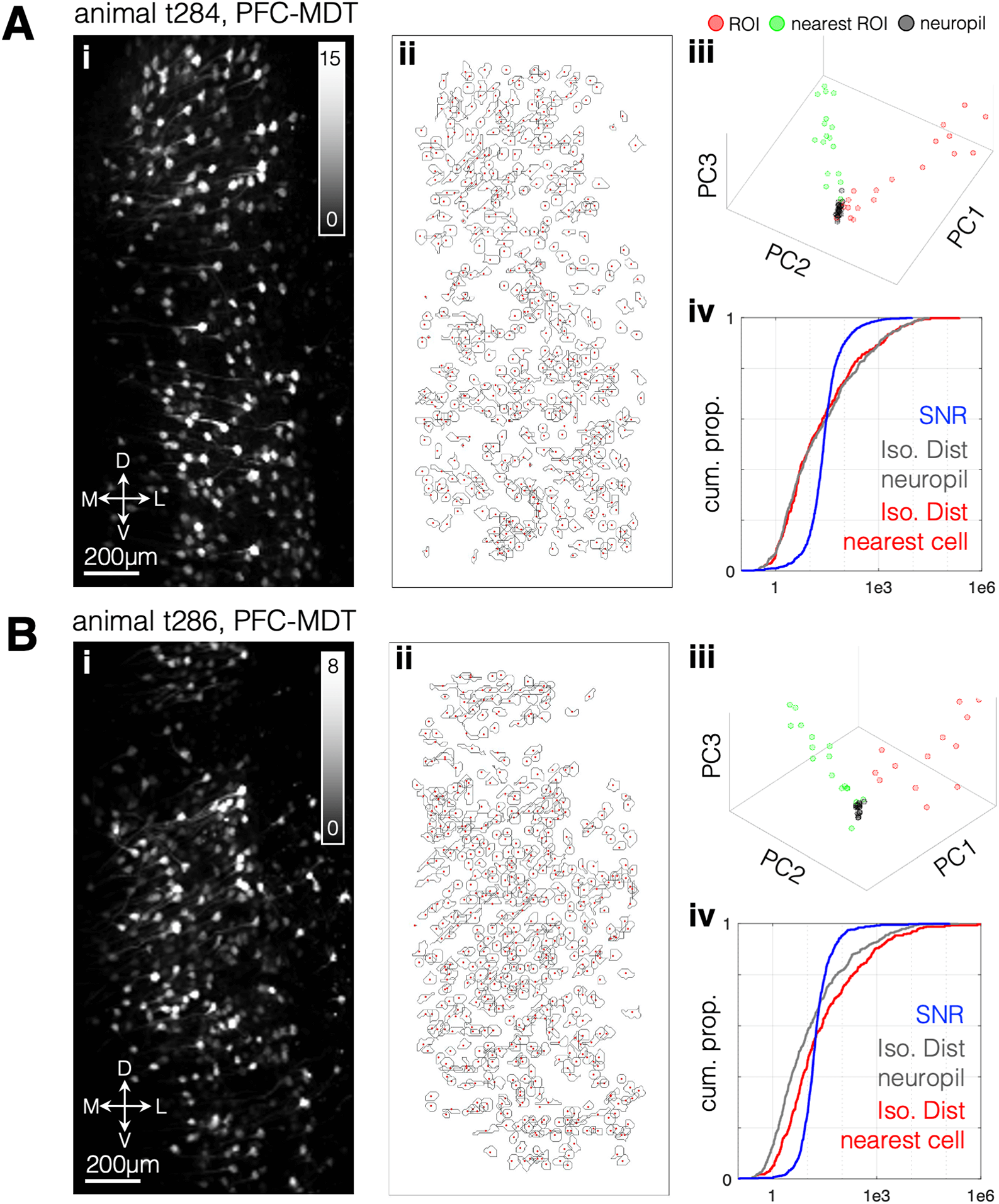
Example Source Localization with CNMF-E, PFC-MDT. A. Left: Session-wide activity projection for an animal labeled with GCaMP6f in PFC-MDT projection neurons. The value for each pixel corresponds with the pixel’s standard deviation across all frames of imaging, normalized to its mean value (its session z-score). Center: Putative cell footprints extracted from the recording using CNMF-E. Grey contours are boundaries of spatial components, red dots are median centroids (N=501 putative cells, 15437 frames, 0.289 f/s, galvano scanning). iii: First three PCA scores for each pixel in a represenative putative cell, its nearest neighbor, and neuropil, defined as an equal or greater number of pixels drawn from immediately outside the footprint and not within any other footprint. iv: Population cumulative distributions of isolation distances for the population. Red: isolation from nearest neighbor. Gray: isolation from neuropil. Blue: signal/noise ratios for pixels from identified neurons. B. Same as (A), for a second PFC-MDT animal.

**Figure S4.**
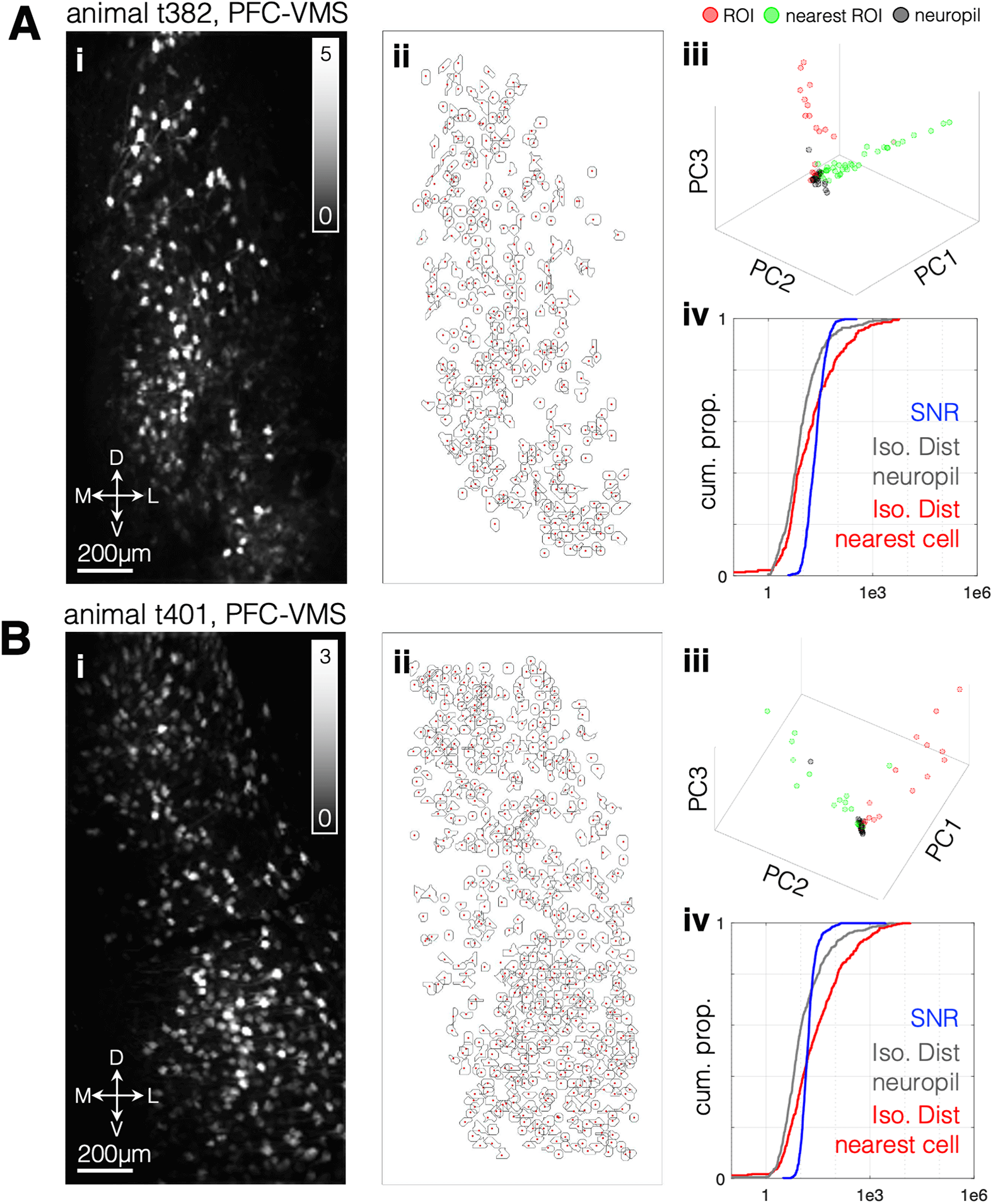
Example Source Localization with CNMF-E, PFC-VMS. Same as Fig. S3 for two PFC-VMS animals.

**Figure S5.**
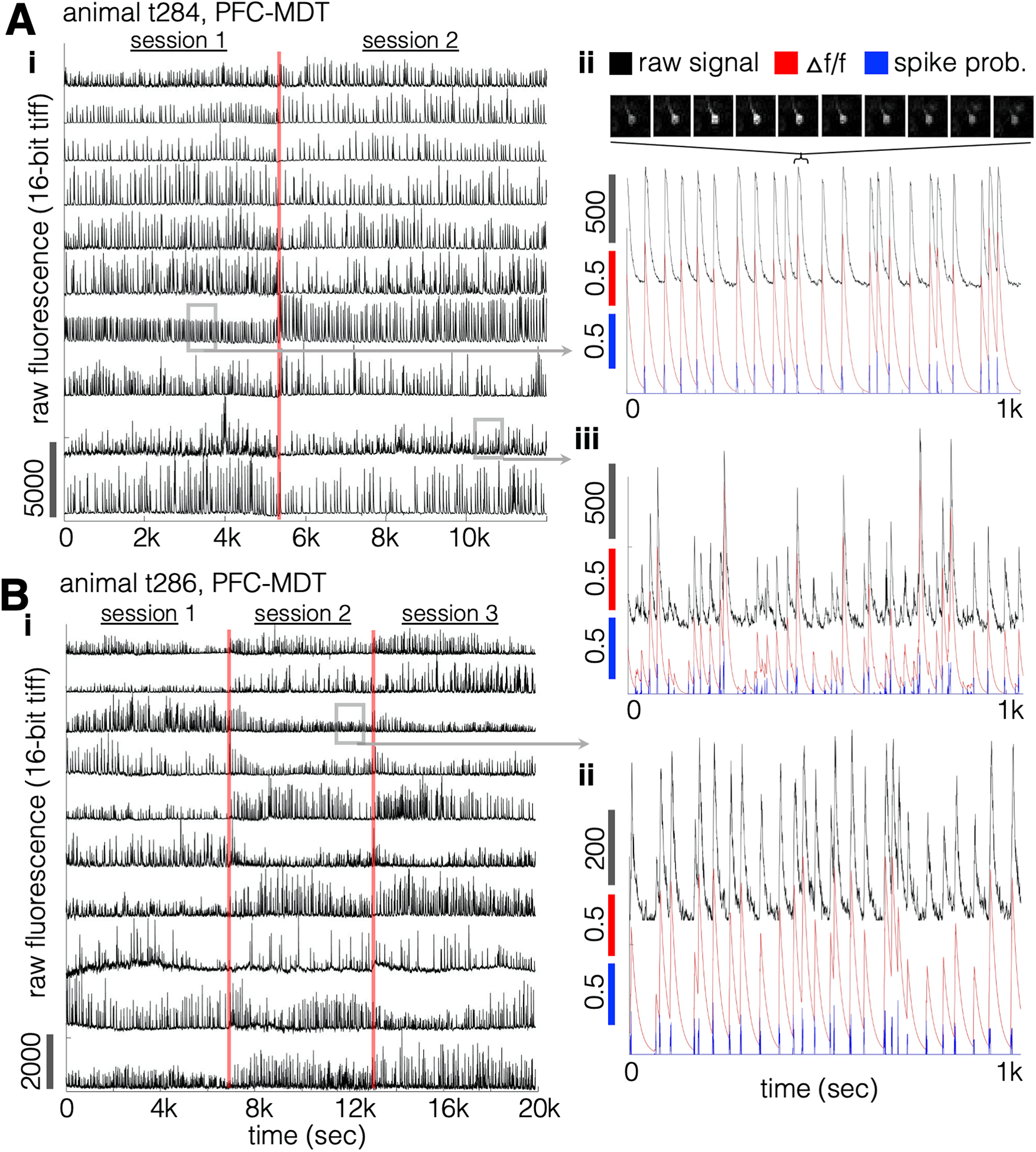
Example Signal Extraction for a PFC-MDT animal. A. Example signals, PFC-MDT, same animal as (S3-A). i) Ten representative raw fluorescence traces from putative neurons identified in two successive SEDS sessions. ii) (top): Individual video frames from a single calcium transient event. Frames shown are downsampled to a rate of ½ the recording rate. Bottom: traces from the corresponding neuron as raw fluorescence (black, units are pixel brightness as unsigned 16-bit integers, maximum value 65,536), denoised △f/f (red), and deconvolved instantaneous spike pobability (blue). B. As in (A), for a second PFC-MDT animal (same as S3-B).

**Figure S6.**
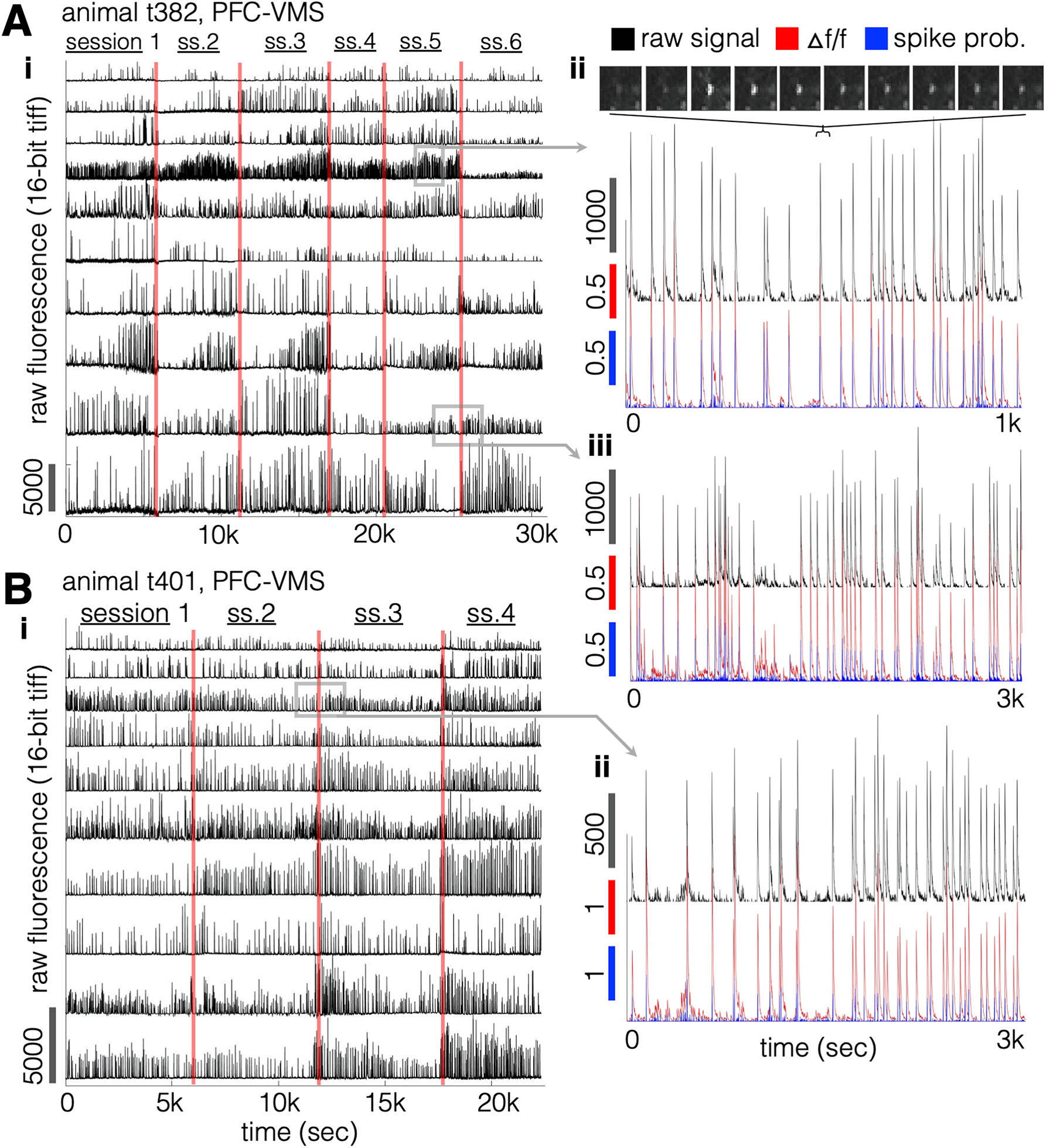
Additional examples of signal extraction. As in S5. Same animals as in S4.

**Figure S7.**
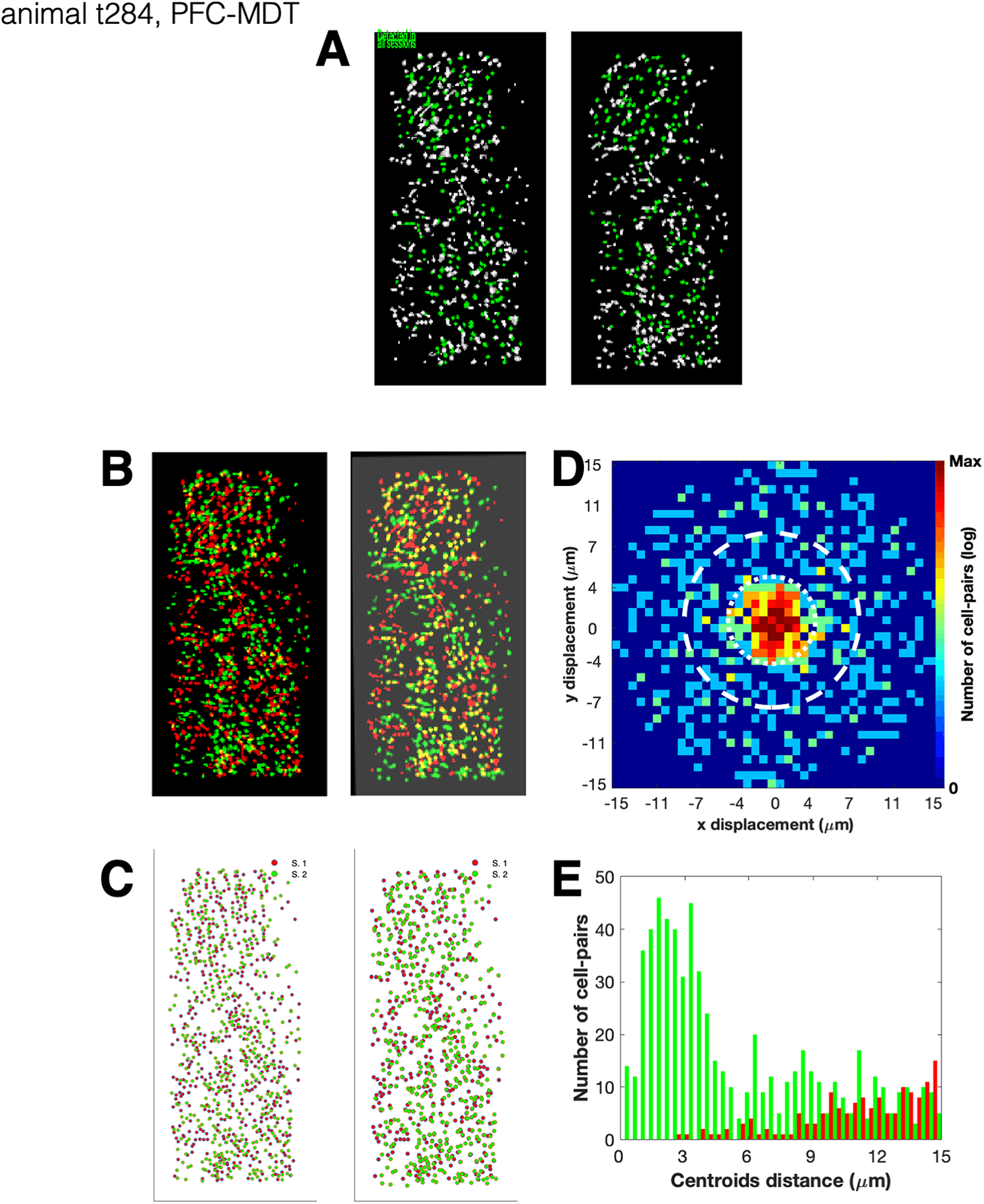
Example cross-session cell footprint alignment using the CellReg cell registration algorithm. A. CNMF-E-extracted cell footprints from session 1 and 2, same animal as in S3-A and S5-A. Green-filled footprints are those detected across all recording sessions. B. Pre-aligned (left) and post-aligned (right) cell footprints. Non-rigid alignment method. C. Pre-aligned (left) and post-aligned (right) cell centroids. D. Post-alignment offset of putative matching cell footprints. E. Centroid distance histograms for putative matching (green) and non-matching (red) pairs of footprints.

**Figure S8.**
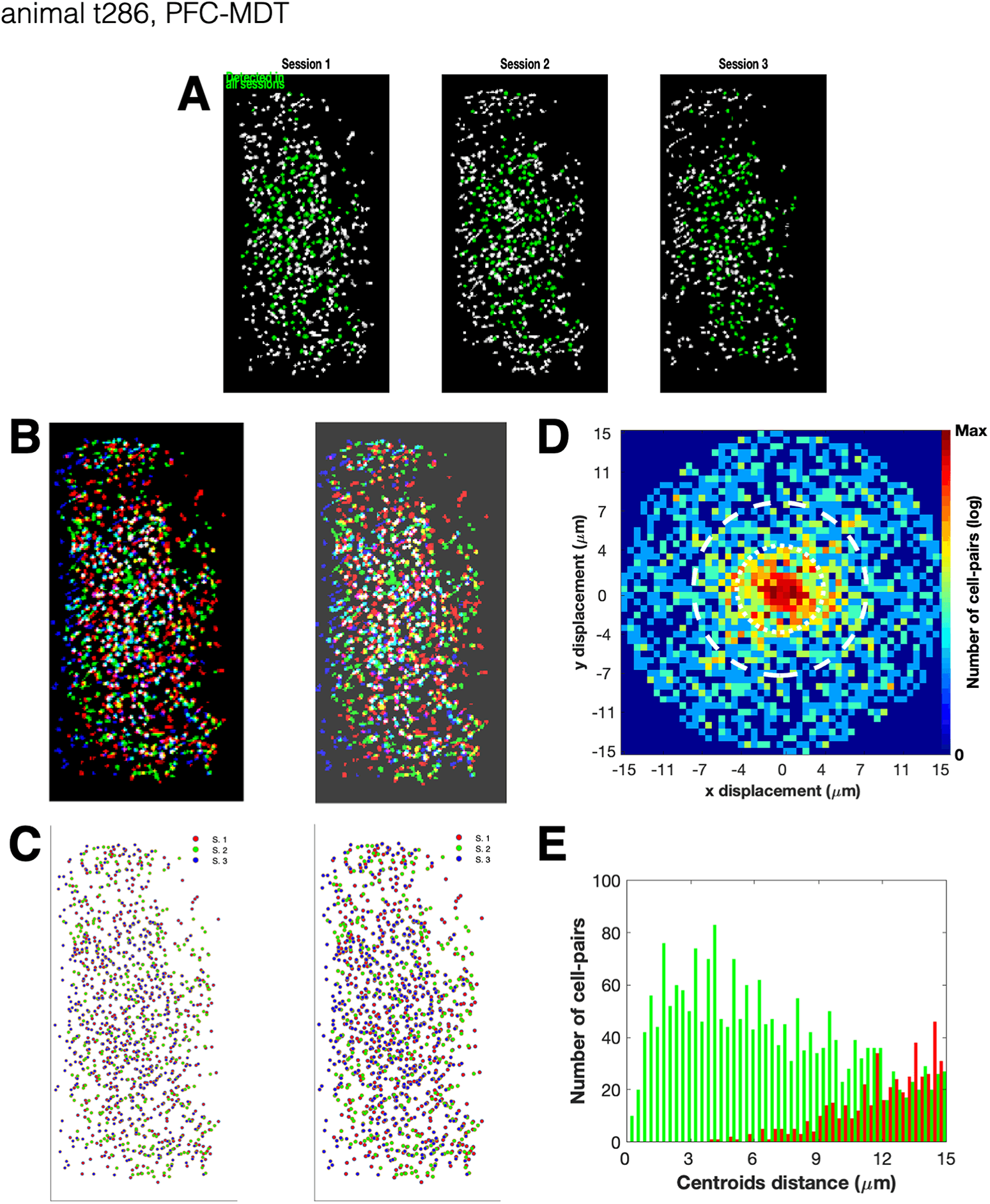
Additional examples of cross-session cell footprint alignments. Animals from S3-B, S4-A, and S4-B, respectively.

**Figure S9.**
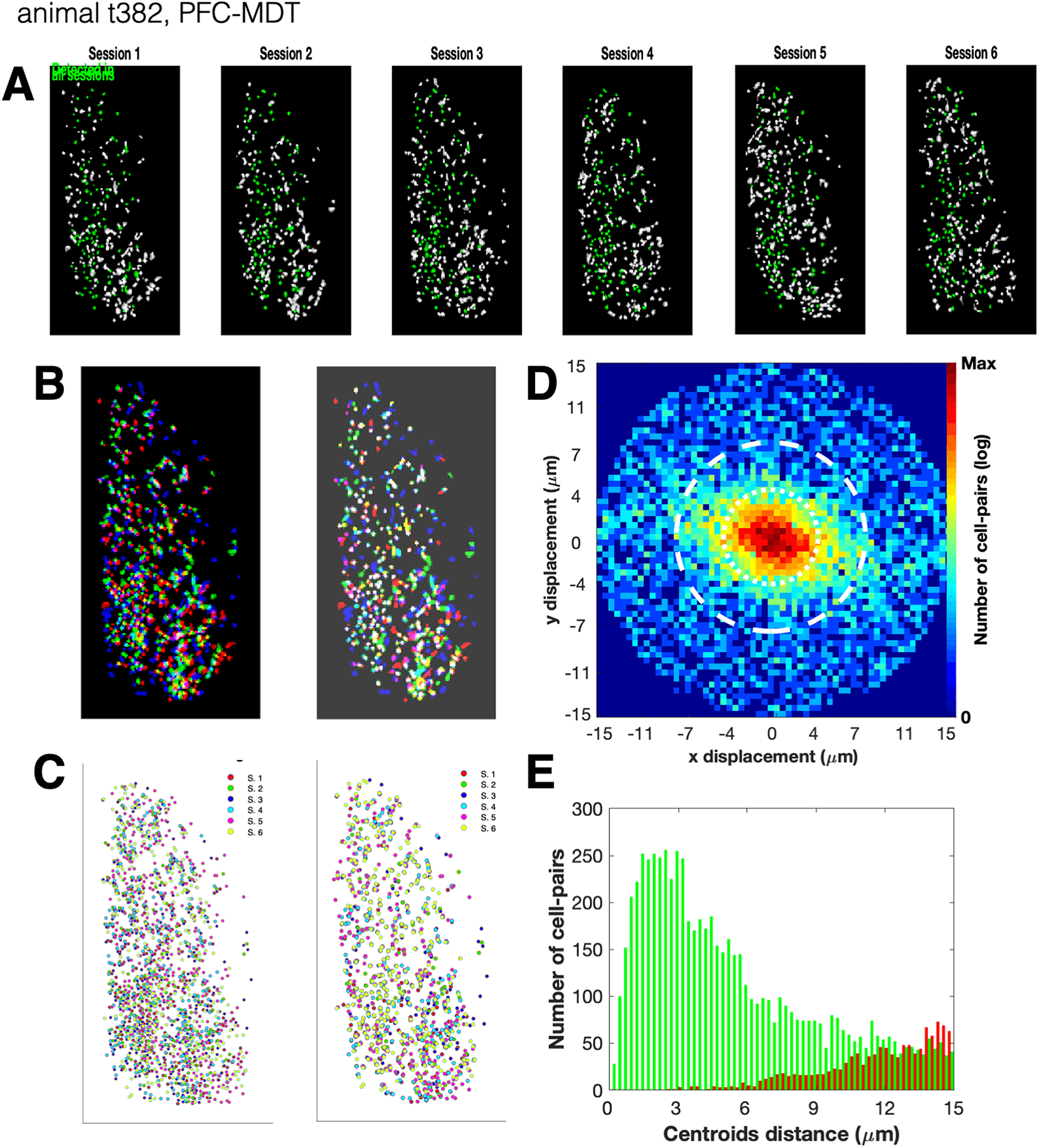
Additional examples of cross-session cell footprint alignments. Animals from S3-B, S4-A, and S4-B, respectively.

**Figure S10.**
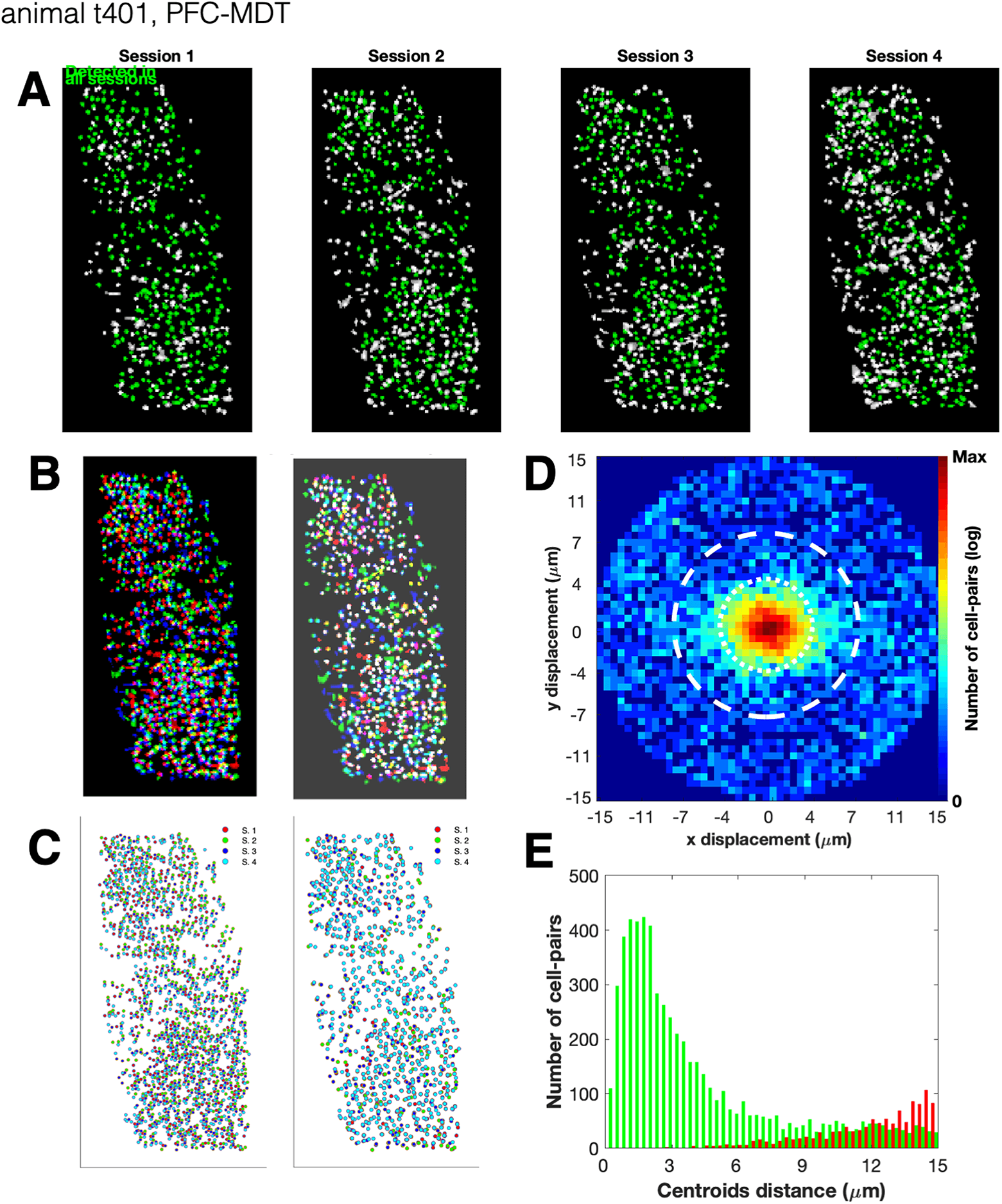
Additional examples of cross-session cell footprint alignments. Animals from S3-B, S4-A, and S4-B, respectively.

**Figure S11.**
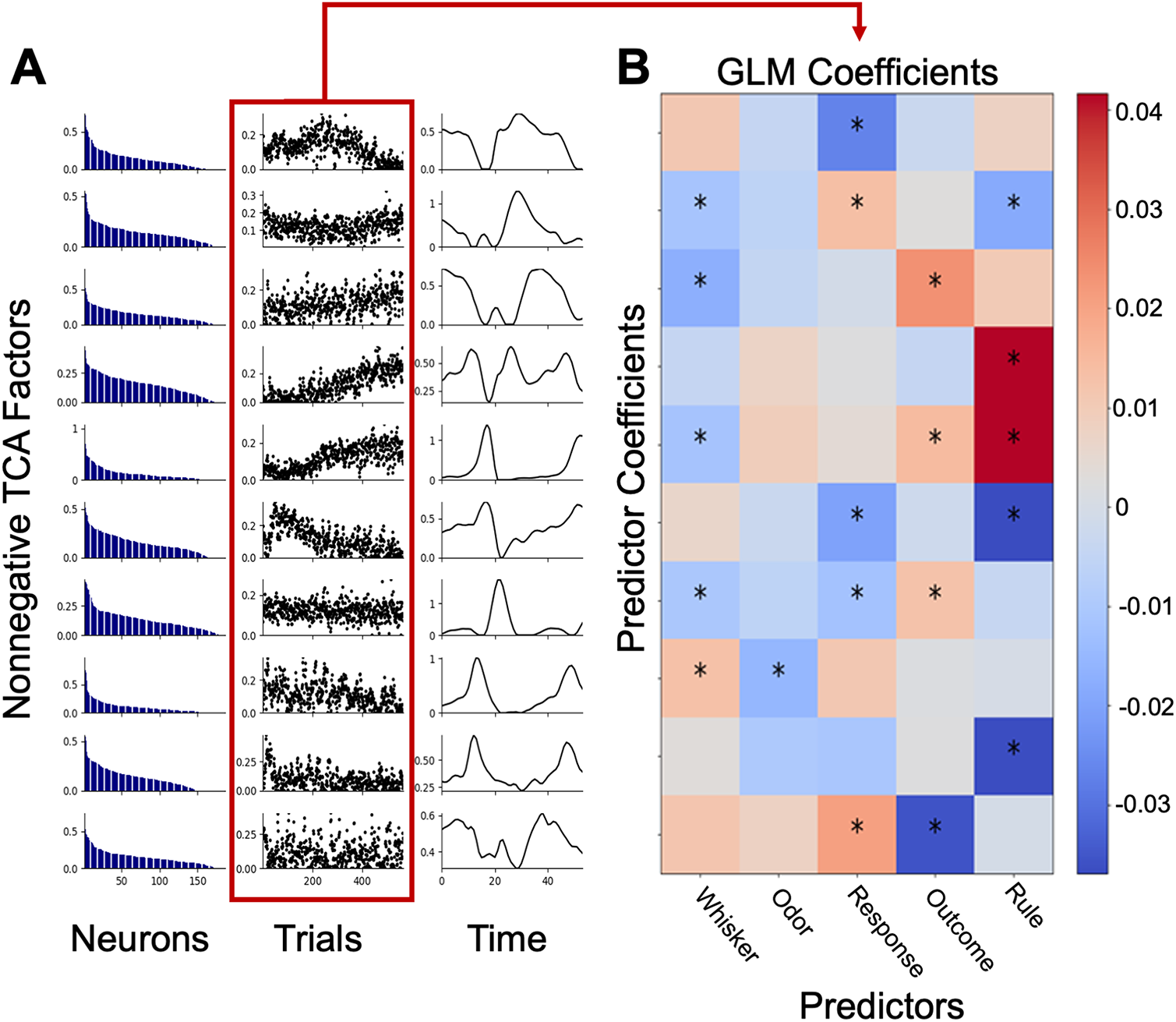
Task modulation of nonnegative TCA Factors. A. Representative factor estimates identified using nonnegative TCA as defined by neuron, trial, and time components. Data shown is from a single mouse and session (186 neurons × 566 trials × 54 timepoints) decomposed into 10 factors. Neuron components are sorted for each factor independently. B. Trial components for each factor were then regressed using a GLM onto task variables. Asterisks denote significance (t-statistic, p<0.05).

**Figure S12.**
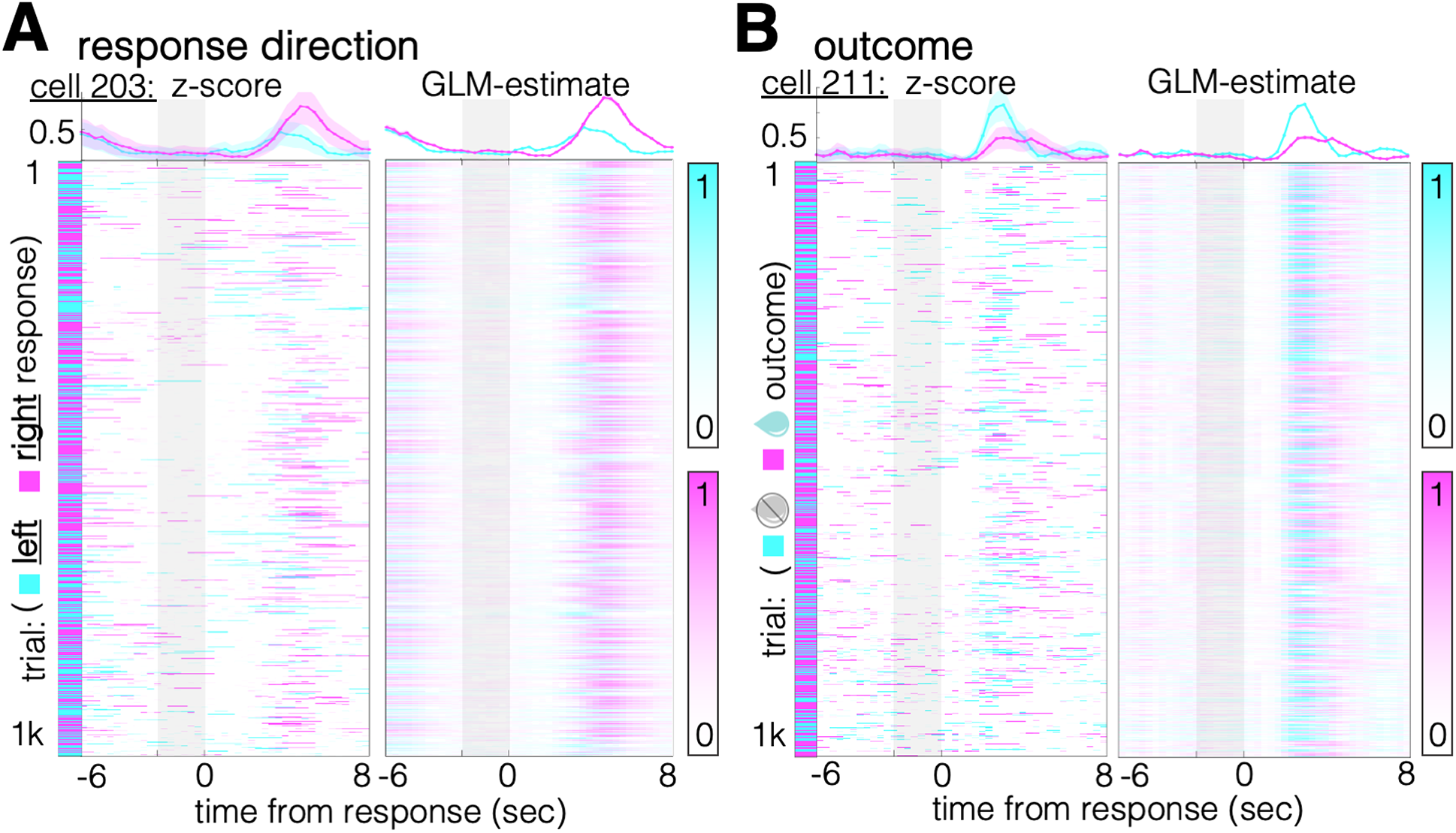
Temporal dynamics of response direction and trial outcome representation. Two additional examples of neurons individually representing response direction and trial outcome, accompanying the examples shown in Fig. 3A. A. Left: z-scored activity on 1071 individual trials (y-axis), and mean z-scored activity (top) over the pre-trial, trial, and post-trial ITI epochs (x-axis) for trials on which the animal responded left (cyan) and for which the animal responded right (magenta). Right: GLM estimate of trial-specific activity. B. Left: activity rastors (y-axis) and mean activity (top) for a second neuron, same session. Cyan: incorrect trials; magenta: correct trials. Right: GLM estimate of trial-specific activity.

**Figure S13.**
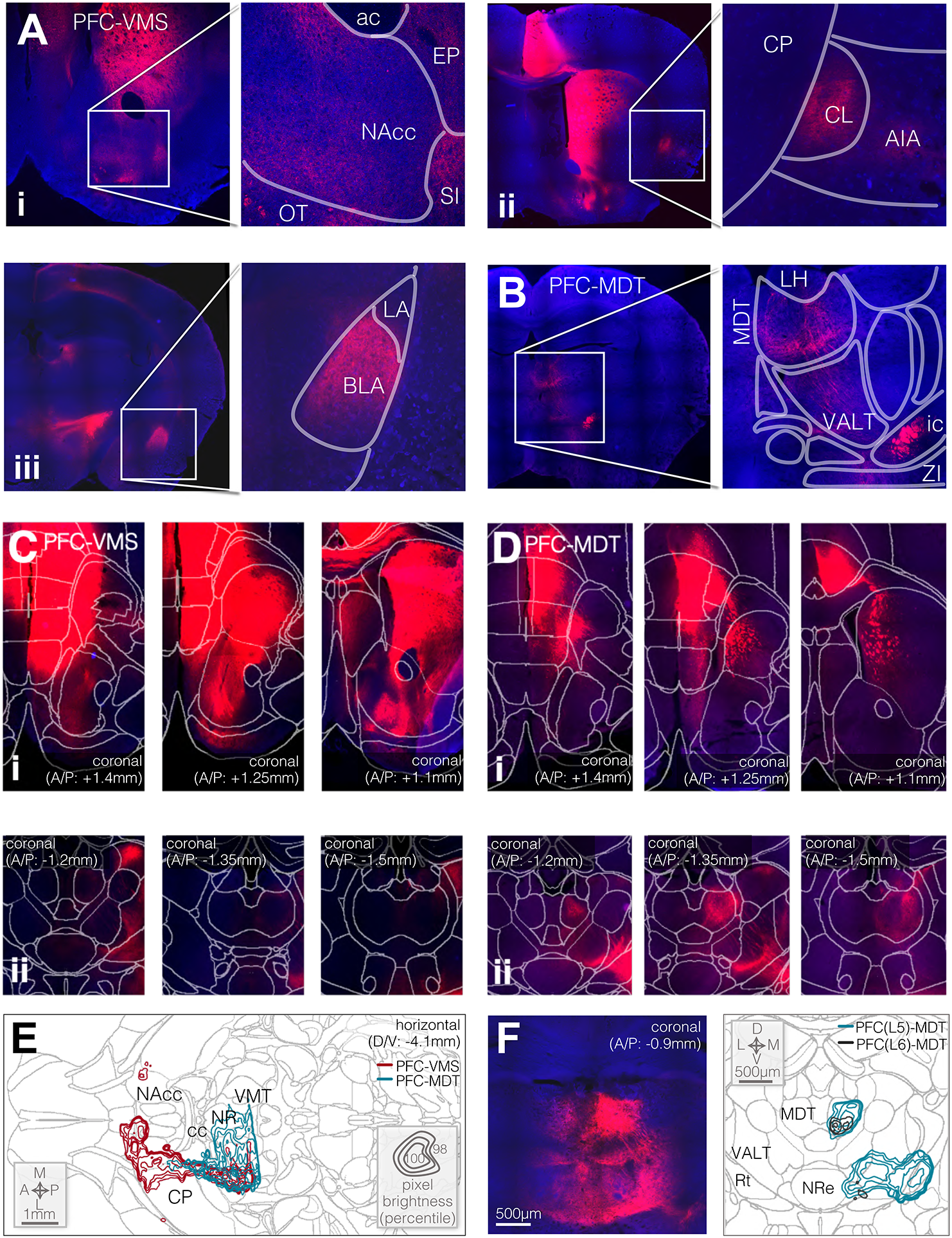
Fluorescence in long-range axons from PFC-VMS and PFC-MDT projection neurons. A. Long-range efferent axons from PFC-VMS neurons. Red: CAG-DIO-tdTomato; blue: DAPI. i) Fluorescent axons in the nucleus accumbens (NAcc), substantia innominata (SI), endopiriform nucleus (EP), and olfactory tubercle (OT). ii) Axon collaterals from PFC-VMS neurons in the claustrum (CL) and agranular insular cortex (AIA). iii) Axon collaterals from PFC-VMS neurons in basolateral amygdala (BLA). B. Fluorescent axons from PFC-MDT neurons in mediodorsal thalamus (MDT), zona incerta (ZI), and the ventral anterolateral thalamic nucleus (VALT). C. Coronal slices at multiple A/P cross-sections from within the same PFC-VMS labeled animal. i) Nucleus accumbens and surrounding regions at +1.4mm, +1.25mm, and +1.1mm relative to bregma (Allen CCF). ii) Medial thalamus and surrounding regions in the same animal, same laser and confocal microscope settings. D. Coronal slices at multiple A/P cross-sections from within the same PFC-MDT labeled animal. i) NAcc and surrounding regions at +1.4mm, +1.25mm, and +1.1mm relative to bregma. ii). Medial thalamus and surrounding regions in the same animal. E. Contour plots of axon fluorescence density from a composite of 5 PFC-VMS and 5 PFC-MDT labeled animals. Horizontal cross-section at 4.1mm ventral to bregma (Allen CCF). Concentric contours correspond with pixel brightness, which is represented as a percentile from each animal’s pixel value distribution. Outermost of five concentric contours corresponds with the 98^th^ percentile, and the innermost enclosed contour corresponding with the 100^th^ percentile for pixel brightness. F. Left: Example coronal cross-section of medial thalamus and surrounding regions from a PFC(L6)-MDT labeled animal (Cav-Cre in MDT and Flex-tdTomato in PFC). right: Composite contour plots for 5 PFC(L5)-MDT (green) and 2 PFC(L6)-MDT (black) labeled animals. Pixel brightness scale same as (E).

**Figure S14.**
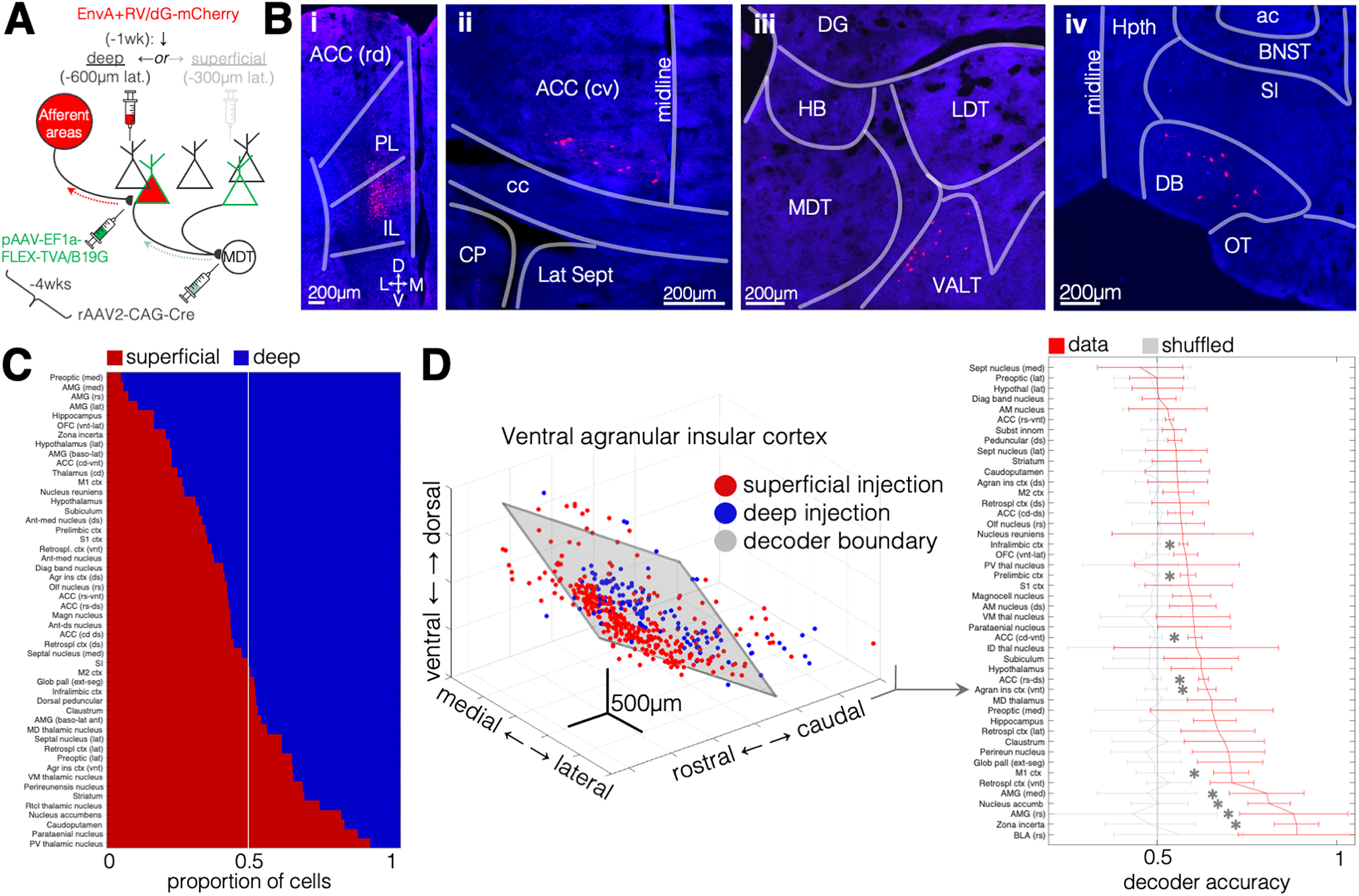
Rabies retrograde labeling experiment. A. Schematic diagram of G-deleted rabies tracing experiment. At 4 weeks prior to sacking, an AAV encoding the TVA rabies B19 glycoprotein (pAAV-EF1a-FLEX-TVA/B19) was injected into PFC, and in the same surgery, retrogradely transported rAAV2-CAG-Cre was injected into MDT or VMS (see Methods). at 1 week prior to sacking, animals underwent an injection of EnvA G-Deleted Rabies-mCherry into either superficial (300um from midline) or deep (600um from midline) PFC, at a volume of 50nL). This approach allows for monosynaptic retrograde labeling of projection neurons. B. Example histological images of EnvA G-Deleted Rabies-mCherry labeling. i: labeling around the injection site. Injections were targeted to the PL/IL border. ii: labeling in caudo-ventral ACC. This region showed enhanced labeling in deep-injected animals relative to superficially-injected animals. iii: labeling in the ventral antero-lateral nucleus of the thalamus, one of numerous thalamic nuclei to show robust labeling, including medio-dorsal thalamus, retrospenial nucleus, paraventricular thalamus, nucleis of reuniens, and parataenial nucleus. iv: labeling in the diagonal band nucleus. Robust labeling was seen throughout the cholinergic nuclei of the basal forebrain, including the medial septum, the preoptic areas, and substantia innominata. C. Relative cell densities in deep vs superficially-injected animals across brain areas. Regions shown are the 50 most cell-rich regions. D. Support-vector-machine-based decoders for within-region classification of cells from the two PFC injection depths. Inputs are the x,y,and z coordinates of each neuron, and each classifier was tested using 20-fold cross-validation and assuming uniform between-class distribution. Left: optimal support vector for ventral agranular insular cortex, which did not show a difference in total cell number between injection types, but in which a rostro-dorsal / caudo-ventral boundary between cell types is observed. Right: decoder performance, as well as performance on shuffled data, for each of the regions shown in C. Error bars: standard deviations. Asterisks: 2-sided t-test, p<0.05.

**Figure S15.**
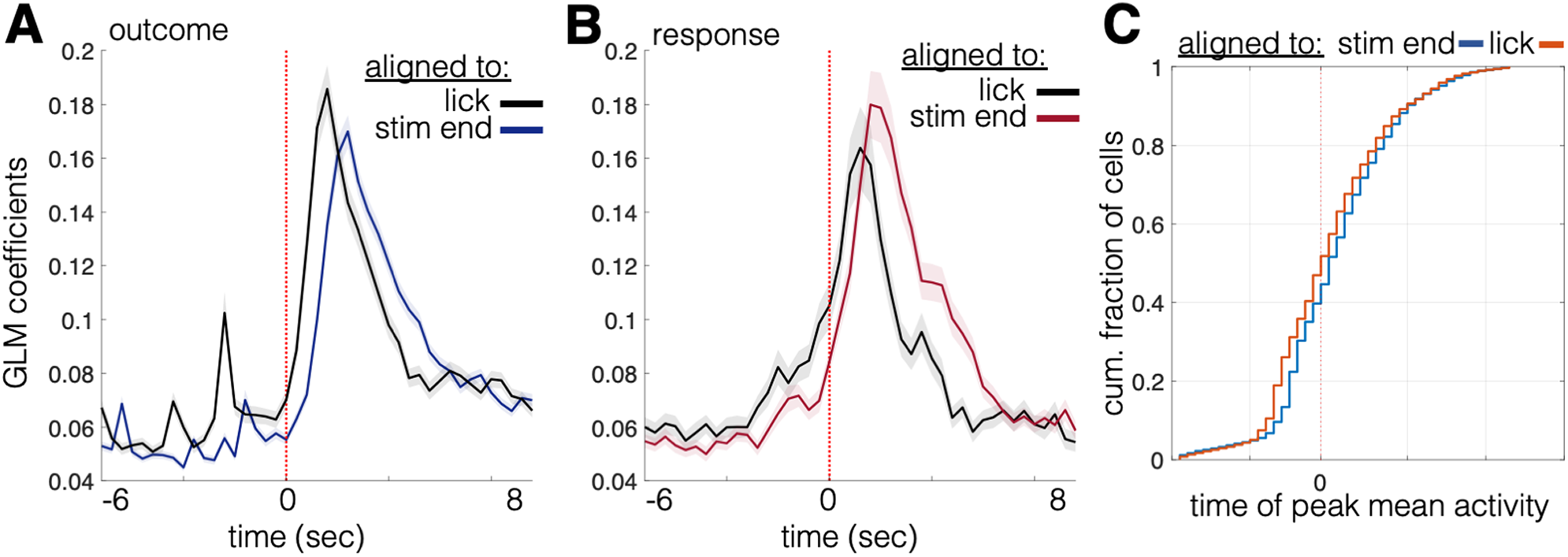
Comparison of response and outcome signals by temporal trial alignment. A. Mean +/− SEM for trial-aligned GLM coefficients of neurons significantly modulated by trial outcome direction. Modulated neurons include those with significance at any timepoint (Bonferroni-corrected for the 43 trial timepoints). Blue: coefficients aligned to stimulus offset; black: coefficients aligned to first lick within the response window. B. Same as (A), for response direction. Red: coefficients aligned to stimulus offset; black: coefficients aligned to first lick within the response window. C. Cumulative histogram of times of peak absolute activity (relative to the mean over the 43 trial timepoints) for each cell. Blue: peaks aligned to stimulus offset; red: peaks aligned to first lick within response window.

**Table S16. Summary of behavioral statistics for optogenetic experiment**

## References

Adesnik, H., and Naka, A. (2018). Cracking the Function of Layers in the Sensory Cortex. Neuron VOLUME 100, ISSUE 5, P1028–1043.

Akrami, A., Kopec, C., Diamond, M., and Brody, C. (2018). Posterior parietal cortex represents sensory history and mediates its effects on behaviour. Nature Feb 15;554(7692):368–372.

Andermann, M., Gilfoy, N., Goldey, G., Sachdev, R., Wölfel, M., McCormick, D., Reid, R., and Levene, M. (2013). Chronic Cellular Imaging of Entire Cortical Columns in Awake Mice Using Microprisms. Neuron 80, 900–913.

Aoki, S., Liu, A.W., Zucca, A., Zucca, S., and Wickens, J.R. (2015). Role of Striatal Cholinergic Interneurons in Set-Shifting in the Rat. J. Neurosci. 35, 9424–9431.

Bari, B., Grossman, C., Lubin, E., Rajagopalan, A., Cressy, J., and Cohen, J. (2019). Stable Representations of Decision Variables for Flexible Behavior. Neuron 103, 922–933.

Bernacchia, A., Seo, H., Lee, D., and Wang, X. (2011). A reservoir of time constants for memory traces in cortical neurons. Nat Neurosci. Mar;14(3):366–72.

Biró, S., Lasztóczi, B., and Klausberger, T. (2019). A Visual Two-Choice Rule-Switch Task for Head-Fixed Mice. Front Behav. Neurosci. 06 June.

Birrell, J., and Brown, V. (2000). Medial frontal cortex mediates perceptual attentional set shifting in the rat. J. Neurosci. 20, 4320–4324.

Bissonette, G., and Roesch, M. (2015). Neural correlates of rules and conflict in medial prefrontal cortex during decision and feedback epochs. Front Behav. Neurosci. 9: 266.

Bissonette, G., Martins, G., Franz, T., Harper, E., Schoenbaum, G., and Powell, E. (2008). Double dissociation of the effects of medial and orbital prefrontal lesions on attentional and affective shifts in mice. J. Neurosci. 28, 11124–11130.

Bissonette, G., Powell, E., and Roesch, M. (2013). Neural structures underlying set-shifting: roles of medial prefrontal cortex and anterior cingulate cortex. Behav. Brain Res. 250, 91–101.

Block, A., Dhanji, H., Thompson-Tardif, S., and Floresco, S. (2007). Thalamic-prefrontal cortical-ventral striatal circuitry mediates dissociable components of strategy set shifting. Cereb. Cortex 17.

Bortolato, B., Miskowiak, K., Köhler, C., Maes, M., Fernandez, B., Berk, M., and Carvalho, A. (2016). Cognitive remission: a novel objective for the treatment of major depression? BMC Med. 14, Article number: 9.

Brigman, J., Bussey, T., Saksida, L., and Rothblat, L. (2005). Discrimination of multidimensional visual stimuli by mice: intra- and extradimensional set shifts. Behav. Neurosci. 119, 839–842.

Ceaser, A., Goldberg, T., Egan, M., McMahon, R., Weinberger, D., and Gold, J. (2008). Set-shifting ability and schizophrenia: a marker of clinical illness or an intermediate phenotype? Biol Psychiatry 64, 782–788.

Chen, T., Wardill, T., Sun, Y., Pulver, S., Renninger, S., Baohan, A., Schreiter, E., Kerr, R., Orger, M., Jayaraman, V., et al. (2013). Ultrasensitive fluorescent proteins for imaging neuronal activity. Nature ul 18;499(7458):295–300.

Christianini, N., and Shawe-Taylor, J. (2000). An Introduction to Support Vector Machines and Other Kernel-Based Learning Methods (Cambridge, UK: Cambridge University Press).

Collins, D., Anastasiades, P., Marlin, J., and Carter, A. (2018). Reciprocal Circuits Linking the Prefrontal Cortex with Dorsal and Ventral Thalamic Nuclei. Neuron 98, 366–379.

Corbetta, M., and Shulman, G. (2002). Control of goal-directed and stimulus-driven attention in the brain. Nat Rev Neurosci Mar;3(3):201–15.

Crowe, S., and Ellis-Davies, G. (2014). Longitudinal in vivo two-photon fluorescence imaging. J. Comp. Neurol. 522:1708–1727.

Denk, W., Strickler, J., and Webb, W. (1990). Two-Photon Laser Scanning Fluorescence Microscopy. Science Vol. 248, No. 4951., 73–76.

Desimone, R., and Duncan, J. (1995). Neural mechanisms of selective visual attention. Annu Rev Neurosci 18:193–222.

Disner, S.G., Beevers, C.G., Haigh, E.A.P., and Beck, A.T. (2011). Neural mechanisms of the cognitive model of depression. Nat. Rev. Neurosci. 12, 467–477.

Durstewitz, D., Vittoz, N., Floresco, S., and Seamans, J. (2010). Abrupt transitions between prefrontal neural ensemble states accompany behavioral transitions during rule learning. Neuron 66, 438–448.

Ellwood, I., Patel, T., Wadia, V., Lee, A., Liptak, A., Bender, K., and Sohal, V. (2017). Tonic or Phasic Stimulation of Dopaminergic Projections to Prefrontal Cortex Causes Mice to Maintain or Deviate from Previously Learned Behavioral Strategies. J Neurosci 37(35): 8315–8329.

Floresco, S., Ghods-Sharifi, S., Vexelman, C., and O, M. (2006). Dissociable Roles for the Nucleus Accumbens Core and Shell in Regulating Set Shifting. J. Neurosci. 26, 2449–2457.

Fox, M., Barense, M., and Baxter, M. (2003). Perceptual attentional set-shifting is impaired in rats with neurotoxic lesions of posterior parietal cortex. J Neurosci Jan 15;23(2):676–81.

Friedrich, J., Zhou, P., and Paninski, L. (2017). Fast online deconvolution of calcium imaging data. PLoS Comput. Biol. https://doi.org/10.1371/journal.pcbi.1005423.

Garrison, J., Erdeniz, B., and Done, J. (2013). Prediction error in reinforcement learning: a meta-analysis of neuroimaging studies. Neurosci Biobehav Rev Aug;37(7):1297–310.

Gonda, X., Pompili, M., Serafini, G., Carvalho, A.F., Rihmer, Z., and Dome, P. (2015). The role of cognitive dysfunction in the symptoms and remission from depression. Ann. Gen. Psychiatry 14, 27.

Halleland, H., Haavik, J., and Lundervold, A. (2012). Set-shifting in adults with ADHD. J Int Neuropsychol Soc 18, 728–737.

Harris, K., and Shepherd, G. (2015). The neocortical circuit: themes and variations. Nat. Neurosci. 18, pages170–181.

Harvey, P., Green, M., Keefe, R., and Velligan, D. (2004). Cognitive Functioning in Schizophrenia: A Consensus Statement on Its Role in the Definition and Evaluation of Effective Treatments for the Illness. J. Clin. Psychiatry 65, 361–372.

Hauser, T., Iannaccone, R., Staempfli, P., and Drechsler, R. (2013). The Feedback-Related Negativity (FRN) revisited: New insights into the localization, meaning and network organization. NeuroImage 84.

Heisler, J.M., Morales, J., Donegan, J.J., Jett, J.D., Redus, L., and O’Connor, J.C. (2015). The Attentional Set Shifting Task: A Measure of Cognitive Flexibility in Mice. J. Vis. Exp. JoVE.

Hill, S.K., Bishop, J.R., Palumbo, D., and Sweeney, J.A. (2010). Effect of second-generation antipsychotics on cognition: current issues and future challenges. Expert Rev. Neurother. 10, 43–57.

Hyman, J., Holroyd, C., and Seamans, J. (2017). A Novel Neural Prediction Error Found in Anterior Cingulate Cortex Ensembles. Neuron Jul 19;95(2):447–456.e3.

Hymana, J., Maa, L., Balaguer-Ballesterb, E., Durstewitzb, D., and Seamansa, J. (2012). Contextual encoding by ensembles of medial prefrontal cortex neurons. PNAS 109, 5086–5091.

Jazbec, S., Pantelis, C., Robbins, T., Weickert, T., Weinberger, D., and Goldberg, T. (2007). Intra-dimensional/extra-dimensional set-shifting performance in schizophrenia: Impact of distractors. Schizophr. Res. 89, 339–349.

Kato, S., Fukabori, R., Nishizawa, K., Okada, K., Yoshioka, N., Sugawara, M., Maejima, Y., Shimomura, K., Okamoto, M., Eifuku, S., et al. (2018). Action Selection and Flexible Switching Controlled by the Intralaminar Thalamic Neurons. Cell Rep. 27;22(9):2370–2382.

Kim, C., Cilles, S.E., Johnson, N.F., and Gold, B.T. (2012). Domain General and Domain Preferential Brain Regions Associated with Different Types of Task Switching: A Meta-Analysis. Hum. Brain Mapp. 33, 130–142.

Liu, X., and Carter, A. (2018). Ventral Hippocampal Inputs Preferentially Drive Corticocortical Neurons in the Infralimbic Prefrontal Cortex. J Neurosci Aug 15;38(33):7351–7363.

Ljubojevic, V., Luu, P., and De Rosa, E. (2014). Cholinergic Contributions to Supramodal Attentional Processes in Rats. J. Neurosci. 34(6):2264–2275.

Low, R.J., Gu, Y., and Tank, D.W. (2014). Cellular resolution optical access to brain regions in fissures: Imaging medial prefrontal cortex and grid cells in entorhinal cortex. Proc. Natl. Acad. Sci. 111, 18739–18744.

Lui, J., Nguyen, N., Grutzner, S., Darmanis, S., Peixoto, D., Wagner, M., Allen, W., Kebschull, J., Richman, E., Ren, J., et al. (2020). Differential encoding in prefrontal cortex projection neuron classes across cognitive tasks. Cell December 17, 2020 DOI:https://doi.org/10.1016/j.cell.2020.11.046.

MacDonald, A. 3rd, Cohen, J., Stenger, V., and Carter, C. (2000). Dissociating the role of the dorsolateral prefrontal and anterior cingulate cortex in cognitive control. Science.

Mahn, M., Gibor, L., Patil, P., Malina, K., Oring, S., Printz, Y., Levy, R., Lampl, I., and Yizhar, O. (2018). High-efficiency optogenetic silencing with somatargeted anion-conducting channelrhodopsins. Nat. Commun. 9:4125.

Mante, V., Sussillo, D., Shenoy, K., and Newsome, W. (2013). Context-dependent computation by recurrent dynamics in prefrontal cortex. Nature 7;503(7474):78–84.

Marshel, J., KIm, Y., Machado, T., Quirin, S., Benson, B., Kadmon, J., Raja, C., Chibukhchyan, A., Ramakrishnan, C., Inoue, M., et al. (2019). Cortical layer-specific critical dynamics triggering perception. Science Aug 9;365(6453):eaaw5202.

Marton, T., Seifikar, H., Luongo, F., Lee, A., and Sohal, V. (2018). Roles of Prefrontal Cortex and Mediodorsal Thalamus in Task Engagement and Behavioral Flexibility. J Neurosci 38(10):2569–2578.

Meyers, E., Freedman, D., Kreiman, G., Miller, E., and Poggio, T. (2008). Dynamic population coding of category information in inferior temporal and prefrontal cortex. J Neurophysiol 100, 1407–1419.

Miller, P. (2016). Dynamical systems, attractors, and neural circuits. F1000Research 5.

Miller, E., and Cohen, J. (2001). An integrative theory of prefrontal cortex function. Annu Rev Neurosci 24:167–202.

Milner, B. (1963). Effects of different brain lesions on card sorting: The role of the frontal lobes. Arch. Neurol. 9, 100–110.

Murphy, F.C., Michael, A., and Sahakian, B.J. (2012). Emotion modulates cognitive flexibility in patients with major depression. Psychol. Med. 42, 1373–1382.

Nakai, J., Ohkura, M., and Imoto, K. (2001). A high signal-to-noise Ca2+ probe composed of a single green fluorescent protein. Nat. Biotechnol. 19;137–141.

Nakayama, H., Ibañez-Tallon, I., and Heintz, N. (2018). Cell-type specific contributions of medial prefrontal neurons to flexible behaviors. J Neurosci. 10.1523/JNEUROSCI.3537-17.2018.

Otis, J.M., Namboodiri, V.M.K., Matan, A.M., Voets, E.S., Mohorn, E.P., Kosyk, O., McHenry, J.A., Robinson, J.E., Resendez, S.L., Rossi, M.A., et al. (2017). Prefrontal cortex output circuits guide reward seeking through divergent cue encoding. Nature 543, 103–107.

Pnevmatikakis, E., and Giovannucci, A. (2017). NoRMCorre: An online algorithm for piecewise rigid motion correction of calcium imaging data. J. Neurosci. Methods Volume 291, Pages 83–94.

Pnevmatikakis, E.A., Soudry, D., Gao, Y., Machado, T.A., Merel, J., Pfau, D., Reardon, T., Mu, Y., Lacefield, C., Yang, W., et al. (2016). Simultaneous Denoising, Deconvolution, and Demixing of Calcium Imaging Data. Neuron 89, 285–299.

Prado, V., Janickova, H., Al-Onaizi, M., and Prado, M. (2017). Cholinergic circuits in cognitive flexibility. Neuroscience Mar 14;345:130–141.

Proulx, E., Piva, M., Tian, M., Bailey, C., and Lambe, E. (2014). Nicotinic acetylcholine receptors in attention circuitry: the role of layer VI neurons of prefrontal cortex. Cell Mol Life Sci 71(7): 1225–1244.

Rich, E., and Shapiro, M. (2007). Prelimbic/Infralimbic Inactivation Impairs Memory for Multiple Task Switches, But Not Flexible Selection of Familiar Tasks. J Neurosci Apr 25; 27(17): 4747–4755.

Rich, E., and Shapiro, M. (2009). Rat prefrontal cortical neurons selectively code strategy switches. J. Neurosci. 29, 7208–7219.

Rigotti, M., Barak, O., Warden, M., Wang, X., Daw, N., Miller, E., and Fusi, S. (2013). The importance of mixed selectivity in complex cognitive tasks. Nature 497, 585–590.

Rodgers, C., and DeWeese, M. (2014). Neural Correlates of Task Switching in Prefrontal Cortex and Primary Auditory Cortex in a Novel Stimulus Selection Task for Rodents. Neuron 82, 1157–1170.

Sayyah, M., Eslami, K., AlaiShehni, S., and Kouti, L. (2016). Cognitive Function before and during Treatment with Selective Serotonin Reuptake Inhibitors in Patients with Depression or Obsessive-Compulsive Disorder. Psychiatry J. 2016.

Schmitt, L., Wimmer, R., Nakajima, M., Happ, M., MofakHam, S., and Halassa, M. (2017). Thalamic amplification of cortical connectivity sustains attentional control. Nature May 11;545(7653):219–223.

Schmitzer-Torbert, N., Jackson, J., Henze, D., Harris, K., and Redish, A. (2005). Quantitative measures of cluster quality for use in extracellular recordings. Neuroscience 131, 1–11.

Senzai, Y., Fernandez-Ruiz, A., and Buzsaki, G. (2019). Layer-Specific Physiological Features and Interlaminar Interactions in the Primary Visual Cortex of the Mouse. Neuron VOLUME 101, ISSUE 3, P500–513.E5.

Shamash, P., Carandini, M., Harris, K., and Steinmetz, N. (2018). A tool for analyzing electrode tracks from slice histology. BioRxiv doi: https://doi.org/10.1101/447995.

Sharif, F., Tayebi, B., Buzsaki, G., Royer, S., and Fernandez-Ruiz, A. (2020). Subcircuits of Deep and Superficial CA1 Place Cells Support Efficient Spatial Coding across Heterogeneous Environments. Neuron Nov 14;S0896-6273(20)30858-8.

Sheintuch, L., Rubin, A., Brande-Eilat, N., Geva, N., Sadeh, N., Pinchasof, O., and Ziv, Y. (2017). Tracking the Same Neurons across Multiple Days in Ca2+ Imaging Data. Cell Rep. 21, 1102–1115.

Siniscalchi, M., Phoumthipphavong, V., Ali, F., Lozano, M., and Kwan, A. (2016). Fast and slow transitions in frontal ensemble activity during flexible sensorimotor behavior. Nat. Neurosci. Sep;19(9):1234–42.

Siniscalchi, M., Wang, H., and Kwan, A. (2019). Enhanced Population Coding for Rewarded Choices in the Medial Frontal Cortex of the Mouse. Cereb Cortex Sep 13;29(10):4090–4106.

Smith, M., and Kohn, A. (2008). Spatial and Temporal Scales of Neuronal Correlation in Primary Visual Cortex. J Neurosci 28(48): 12591–12603.

Smout, C., Tang, M., Garrido, M., and Mattingley, J. (2019). Attention promotes the neural encoding of prediction errors. PLoS Biol 17(2): e2006812.

Soudais, C., Laplace-Builhe, C., Kissa, K., and Kremer, E. (2001). Preferential transduction of neurons by canine adenovirus vectors and their efficient retrograde transport in vivo. FASEB J 15:2283–2285.

Starkweather, C., Gershman, S., and Uchida, N. (2018). The Medial Prefrontal Cortex Shapes Dopamine Reward Prediction Errors under State Uncertainty. Neuron May 2;98(3):616–629.e6

Stringer, C., and Pachitariu, M. (2019). Computational processing of neural recordings from calcium imaging data. Curr Opin Neurobiol 55, 22–31.

Sul, J., Kim, H., Huh, N., Lee, D., and Jung, M. (2010). Distinct roles of rodent orbitofrontal and medial prefrontal cortex in decision making. Neuron May 13; 66(3): 449–460.

Tait, D.S., Chase, E.A., and Brown, V.J. (2014). Attentional set-shifting in rodents: a review of behavioural methods and pharmacological results. Curr. Pharm. Des. 20, 5046–5059.

Tervo, D., Hwang, B., Viswanathan, S., Gaj, T., Lavzin, M., Ritola, K., Lindo, S., Michael, S., Kuleshova, E., Ojala, D., et al. (2016). A Designer AAV Variant Permits Efficient Retrograde Access to Projection Neurons. Neuron Oct 6. pii: S0896-6273(16)30580-3.

Toyomaki, A., Hashimoto, N., Kako, Y., Murohashi, H., and Kusumi, I. (2017). Neural responses to feedback information produced by self-generated or other-generated decision-making and their impairment in schizophrenia. PLoS ONE 2(8): e0183792.

Turner, J., Parrish, J., Hughes, L., Toth, L., and Caspary, D. (2005). Hearing in Laboratory Animals: Strain Differences and Nonauditory Effects of Noise. Comp Med Feb; 55(1): 12–23.

Wang, Q., Ding, S., Li, Y., Royall, J., Feng, D., Lesnar, P., Graddis, N., Naeemi, M., Facer, B., Ho, A., et al. (2020). The Allen Mouse Brain Common Coordinate Framework: A 3D Reference Atlas. Cell May 14;181(4):936–953.e20.

Williams, A., Kim, T., Wang, F., Vyas, S., Ryu, S., and Shenoy, K. (2018). Unsupervised Discovery of Demixed, LowDimensional Neural Dynamics across Multiple Timescales through Tensor Component Analysis. Neuron 98, 1–17.

Wimmer, R., Schmidt, L., Davidson, T., Nakajima, M., Deisseroth, K., and Halassa, M. (2015). Thalamic control of sensory selection in divided attention. Nature Oct 29;526(7575):705–9.

